# A synthetic CRISPR-Cas nuclease with expanded enzymatic activities

**DOI:** 10.1101/2023.11.12.566739

**Authors:** Ylenia Jabalera, Igor Tascón, Sara Samperio, Jorge P. López-Alonso, Monika Gonzalez-Lopez, Ana M. Aransay, Guillermo Abascal-Palacios, Iban Ubarretxena-Belandia, Raul Perez-Jimenez

**Author notes:** These authors contributed equally to the work (Ylenia Jabalera, Igor Tascón). Corresponding authors (I. U-B) and (R.P-J).

## Abstract

Clustered regularly interspaced short palindromic repeats (CRISPR)-associated endonucleases have revolutionized biotechnology for their potential as programmable genome editors. Yet, most natural nucleases and their variants have limitations. Here, we report a fully synthetic CRISPR-associated (Cas) nuclease (α-synCas) designed by Ancestral Sequence Reconstruction (ASR) that displays a set of robust and distinct targeting properties, not found in any other known CRISPR-Cas Class 2 system. We show that α-synCas is a PAMless nuclease able to catalyse RNA-guided, specific cleavage of dsDNA, ssDNA and ssRNA. The synthetic enzyme is also capable of sequence-nonspecific degradation of dsDNA, ssDNA and ssRNA following activation by complementary dsDNA, ssDNA and ssRNA targets. Furthermore, α-synCas exhibits a robust genome editing activity in human cells and bacteria. Cryo-electron microscopy structures of α-synCas ternary and quaternary complexes provide a framework to understand the structural basis for its expanded enzymatic activities. The capability for programmable multimodal targeting of virtually any nucleic acid sequence distinguishes α-synCas as a promising new tool to extend current CRISPR-based technologies.

## Introduction

CRISPR systems provide bacteria and archaea with adaptive immunity against foreign mobile genetic elements^1–4^. These systems rely on the ability of CRISPR-associated (Cas) nucleases to use CRISPR RNA (crRNA) as a guide, with the additional requirement of a protospacer adjacent motif (PAM), to target and degrade foreign DNA/RNA. The adeptness of CRISPR-Cas nucleases at RNA-guided targeting of nucleic acids has catapulted their repurposing as powerful tools for genome editing and diagnostics^5^. Founded on Cas9 from *Streptococcus pyogenes* (SpCas9)^6,7^, the catalogue of CRISPR-Cas nucleases with novel and improved biotechnological functionalities has been expanded by the discovery of variants of the compact CRISPR class 2 system, including orthologs of Cas9, Cas12 and Cas13 nucleases^2,5,8–11^. Despite the successful search of natural CRISPR-Cas nucleases, important limitations persist^9^, and novel CRISPR tools based on molecular engineering have emerged as promising alternatives ^12–15^. We have previously resurrected active CRISPR-Cas nucleases that no longer exist in nature using Ancestral Sequence Reconstruction (ASR)^12^. These proteins, derived from type II-A CRISPR Cas9 orthologs, retraced the evolution of Cas9 up until modern SpCas9 and displayed enzymatic properties, such as PAMless activity and nucleic acid promiscuity, of great interest for biotechnology.

Here, we exploited ASR purely as a protein design technique to engineer a fully functional, non-natural, synthetic CRISPR-Cas nuclease (α-synCas) derived from orthologs of Cas12a ^16,17^. α-synCas displays a relaxed PAM preference and is able to recognize and cleave complementary dsDNA, ssDNA and ssRNA targets. Upon recognition of the complementary targets, α-synCas also exhibits non-specific (collateral) dsDNA-, ssDNA- and ssRNA-nuclease activity Furthermore, α-synCas can process its own crRNA and shows high genome editing activity in human cells and bacteria. Cryo-electron microscopy (cryo-EM) structures of α-synCas-crRNA bound to a dsDNA target (ternary complex), and to both a dsDNA target and dsDNA collateral substrate (quaternary complex), together with the structure of apo α-synCas, provide a framework to understand the expanded activities of this synthetic enzyme, which is able to virtually target any nucleic acid sequence without restriction. α-synCas shows promise as an innovative and unique tool for CRISPR-based biotechnological applications

## Results

### Structure of α-synCas ternary complex

Sixty-three Cas12a sequences from *Bacillota, Pseudomonadota, Planctomycetota, Spirochaetota* and *Bacteroidota* species (Extended Fig. 1) were used to reconstruct ancient forms of Cas12a following a previously described methodology^12^. The last internal node (with an estimated age of 3000 Mya, CI: 2978.5-3030.4 Mya)^18^ was selected for resurrection resulting in an amino acid sequence with 53 % identity relative to *Francisella novicida* Cas12a (FnCas12a) and a posterior probability average of 0.92. This internal node was selected for the design of α-synCas.

The 3.1 Å resolution cryo-EM structure captures a ternary complex of α-synCas bound to crRNA and target dsDNA containing a T-rich PAM (Fig. 1; PDB: 8QWE; Supplementary Figs. 1 and 2; Extended Table 1). Overall α-synCas is well poised for catalysis and displays a bilobed architecture resembling the characteristic oval “sea conch” shape described for Cas12a^19^. The recognition (REC) lobe comprises both REC1 and REC2 domains, and the nuclease (NUC) lobe comprises the PAM-interacting (PI), wedge (WED), RuvC, and Nuc domains, as well as the bridge helix (BH) (Fig. 1a). The structure provides a snapshot of the ternary complex after the cleavage of both strands of the target dsDNA. The crRNA-target DNA heteroduplex (R-loop, Fig. 1b, Extended Fig. 2) threads through the positively charged central channel between the two lobes. The target strand (TS) hybridizes with the crRNA, while the dissociated non-target strand (NTS) traces a straight path through the grove of the DNA nuclease site. There is unambiguous cryo-EM density downstream of the cleaved end of the TS, which we attribute to a post-cleavage product (Fig. 1c-d, 1d and 1e). As in Cas12a, the nuclease active site of α-synCas lies in a pocket at the interface between Nuc and RuvC. The RuvC domain carries the three (D878, E967 and D1216) highly conserved active site residues (Extended Fig. 3). In contrast to what is observed in other Cas12a enzymes, the nuclease site of α-synCas embraces the dissociated NTS through stacking interactions with W1041 and F971 residues.

**Figure 1.**
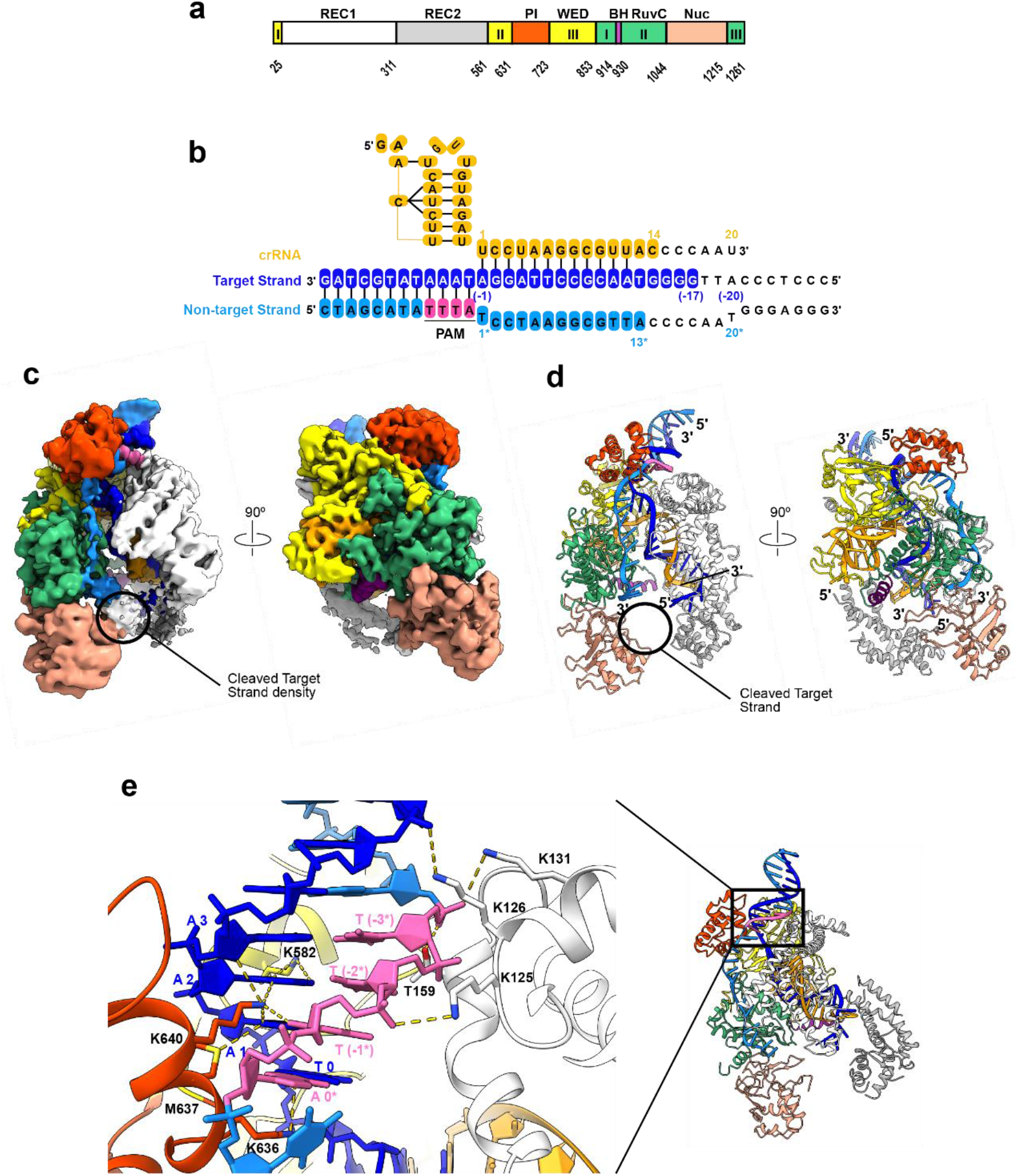
Ternary complex between α-synCas, crRNA and target dsDNA. (**a**) Domain organization of α-synCas. (**b**) Sequences of the crRNA guide and target strand and non-target strand DNA used to form the ternary complex. Nucleotides with coloured background are visible in the cryo-EM map, while uncoloured nucleotides are disordered. (**c**) Unsharpened cryo-EM map of the ternary complex coloured by nucleic acid and protein domain. Unassigned density attributed to the cleaved target strand is highlighted. (**d**) Cryo-EM structure of the a-synCas ternary complex coloured and oriented as in c. A circle marks the putative location of the cleaved target strand. (**e**) Close-up view of the PAM region of the dsDNA highlighting the interactions with selected residues, shown as sticks, of the PI, WED, and REC1 domains.

Cas12a structural studies have suggested a reaction cycle coupled to conformational changes, which begins by an apo state at equilibrium between closed and open conformations, where crRNA shifts the equilibrium to the closed form and target dsDNA causes a further structural tuning that includes the rearrangement of the bridge helix and helix 1 of the RuvC II sub-domain^19–22^. These latter changes in RuvC are absent in the cryo-EM structure of α-synCas ternary complex, likely because the structure represents a post-cleavage form and the enzyme might have reverted to a non-activated state. We note that the 3.3 Å resolution structure of apo α-synCas (Supplementary Note 1, Supplementary Figs. 2 and 3; Extended Table 1) without nucleic acids displays a ring-like architecture distinct from the closed oval “sea conch” shape of the ternary complex (Extended Fig. 4), and from the open and closed conformations suggested for Cas12a in the apo state. The REC2 domain closes the ring by connecting with the RuvC domain, in a way that hinders access to the nuclease active site (Supplementary Fig. 4), and the central channel between the recognition and nuclease lobes, where the crRNA-target DNA heteroduplex threads in the ternary complex, is not formed.

### α-synCas displays targeting flexibility

In contrast to natural CRISPR-Cas effectors, which only exert nuclease activity over one or two types of nucleic acid substrates^11^ (Extended Fig. 5), α-synCas displays a broad *cis*-activity against complementary dsDNA (Fig. 2a), ssDNA (Fig. 2b) and ssRNA (Fig. 2c) targets containing the signature T-rich PAM sequence of canonical Cas12a substrates^16^. *In vitro*, α-synCas/crRNA binary complex cleaves plasmid DNA almost entirely, indicating a robust double-strand break (DSB) activity. The enzyme is also efficient at degrading single-strand substrates, demonstrating that α-synCas behaves as an RNA-guided DNase/RNase. These wide range of activities have only been reported for ancestral Cas9 nucleases^12^.

**Figure 2.**
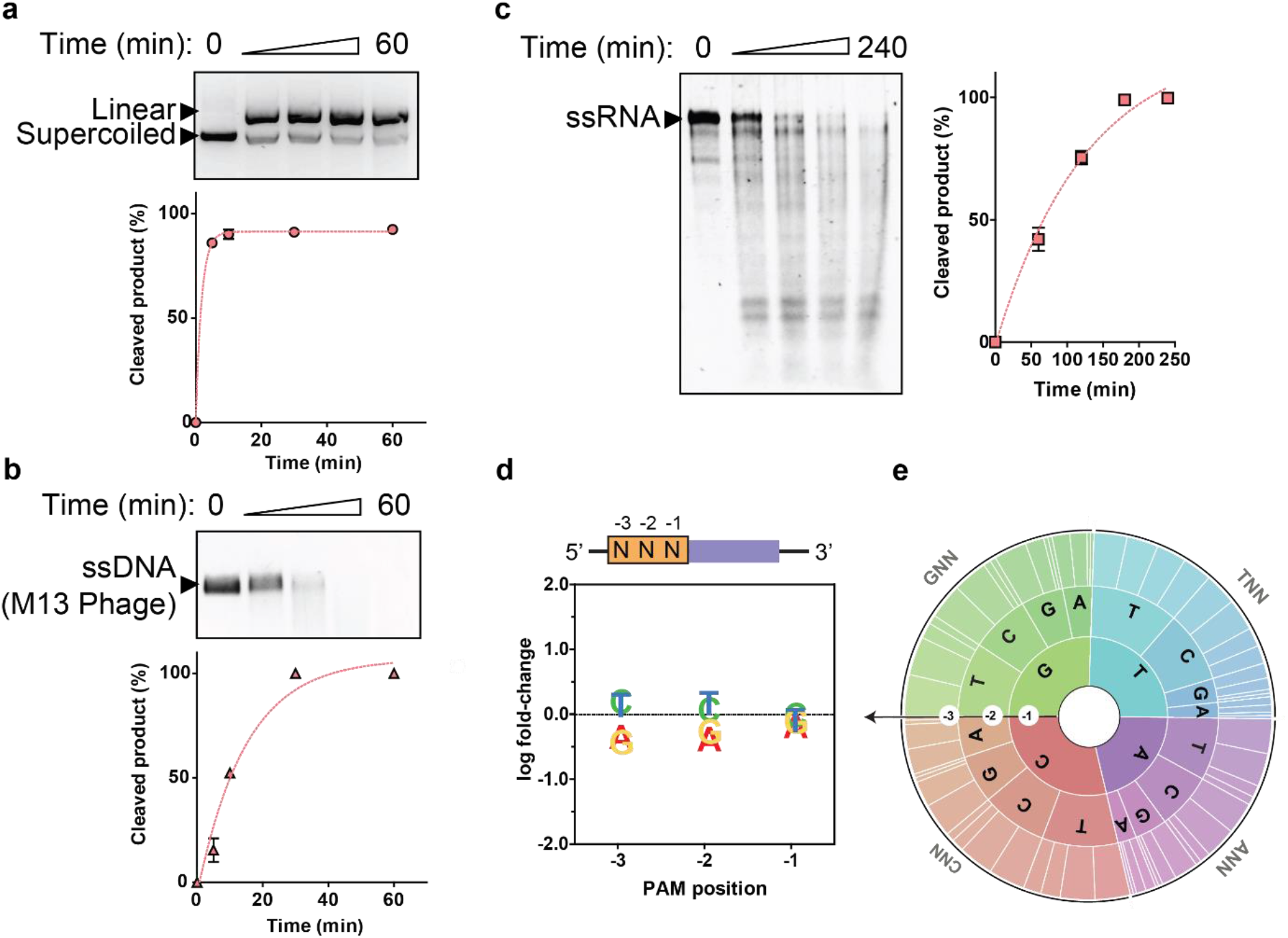
Nuclease activity of α-synCas against dsDNA, ssDNA and ssRNA targets. (**a**) *In vitro* DSB activity of α-synCas against a 4,007 bp supercoiled plasmid DNA as a function of time. Exponential fit of the data expressed as % of cleaved product. (**b**) *In vitro* activity of α-synCas against M13 phage derived ssDNA (M13 phage) as a function of time. Exponential fit of the data expressed as % of cleaved product. (**c**) *In vitro* activity of α-synCas against ssRNA as a function of time. Exponential fit of the data expressed as % of cleaved product. (**d**) PAM wheels (Krona plots) and (**e**) Weblogo with 3 nt PAM recognized by α-synCas. Error bars represent the mean ± SD, where n = 3.

To examine the targeting determinants of α-synCas we measured its PAM and crRNA scaffold recognition requirements. The PAM preference of α-synCas was measured using a DNA library designed with seven random nucleotides (NNNNNNN) followed by a target protospacer. An 855 bp, dsDNA fragment was used as substrate. *In vitro* digestion with α-synCas/crRNA produced a 258 bp-long fragment and a longer 597 bp fragment harbouring the PAM sequence. Next-generation sequencing (NGS) of the longest product revealed a relaxed preference by α-synCas for any PAM tested (Fig. 2d and 2e), suggesting the synthetic nuclease has a minimal PAM requirement. This PAMless activity was verified *in vitro* using non-T-rich PAM sequences (Extended Fig. 6a). Strict PAM recognition is key for dsDNA targeting by CRISPR-Cas nucleases. We have reported that resurrected ancestral forms of *Streptococcus pyogenes* Cas9 display PAM recognition flexibility^12^. Although the structural determinants underlying this flexibility are yet to be determined, we suggested that strict PAM recognition might be the result of evolutionary pressure with the most ancient forms of the enzyme displaying flexible PAM preferences^12^. In the structure of the α-synCas ternary complex both WED and PI domains grip the PAM region of the target dsDNA in a manner analogous to other Cas12a enzymes (Fig. 1e). Uniquely for α-synCas, however, residues K124, K125 and K130 from the REC1 domain stablish electrostatic interactions with the phosphate backbone of the PAM bases through the minor grove (Extended Fig. 7), in a manner that could provide a structural basis for the flexible PAM readout displayed by α-synCas.

To determine the crRNA scaffold recognition determinants of α-synCas, *in vitro* cleavage assays were performed by incubating the enzyme with crRNAs from various CRISPR-system types (Extended Fig. 6b) and Cas12a species (Extended Fig. 6c). α-synCas was able to nick and linearize plasmid DNA using as guides crRNA from different CRISPR-system types. The synthetic enzyme could also linearize plasmid DNA using crRNA from several Cas12a species. This ability to accommodate crRNA scaffolds from the same CRISPR-Cas type has been observed for type V nucleases^23^, however the recognition of non-canonical crRNAs across CRISPR-Cas types has not been reported in any other CRISPR-Cas effector.

We also measured the ability of α-synCas to process its own crRNA (Extended Fig. 6d). The cleavage pattern (65 nt and 49 nt) against a single customized pre-crRNA transcript^24^, corresponding to cuts at the 5’ ends of direct repeat hairpins, was identical to that observed in *E. coli* cells expressing FnCas12a^16^. Thus, α-synCas is able to process its own crRNA guides, similar to type V effector nucleases^25,26^, and can share crRNAs with different types of CRISPR-Cas effectors. This capability of α-synCas should be relevant for multiplexed genome editing applications that employ customized CRISPR arrays for multiple targeting^24^.

### α-synCas exhibits promiscuous collateral activity

Collateral activity (*trans-*nuclease activity) is characteristic of CRISPR Class 2 type V and type VI systems, such as Cas12 and Cas13^17,27–29^, and is defined as the ability to cleavage indiscriminately non-target nucleic acids without complementarity to the crRNA guide. Activation of the nuclease by a target substrate (*cis*-activation) complementary to the crRNA guide is a prerequisite for collateral activity. Upon *cis*-activation of α-synCas by target dsDNA, ssDNA and ssRNA substrates, the synthetic enzyme displayed high degradation activity against dsDNA (Fig. 3a), ssDNA (Fig. 3b) and ssRNA (Fig. 3c) non-target substrates. Thus, α-synCas can efficiently degrade all forms of nucleic acids in *trans* upon activation by different *cis*-activators. This promiscuous collateral activity displayed by α-synCas has not been reported for natural CRISPR-Cas effectors. Indeed, other type V and VI Cas effectors indiscriminately degrade ssDNA only after *cis*-activation by dsDNA targeting (Cas12a)^17,27^, ssRNA after *cis*-activation by ssRNA (Cas13)^28^, or dsDNA, ssDNA and ssRNA after *cis*-activation by ssRNA (Cas12a2)^29^.

**Figure 3.**
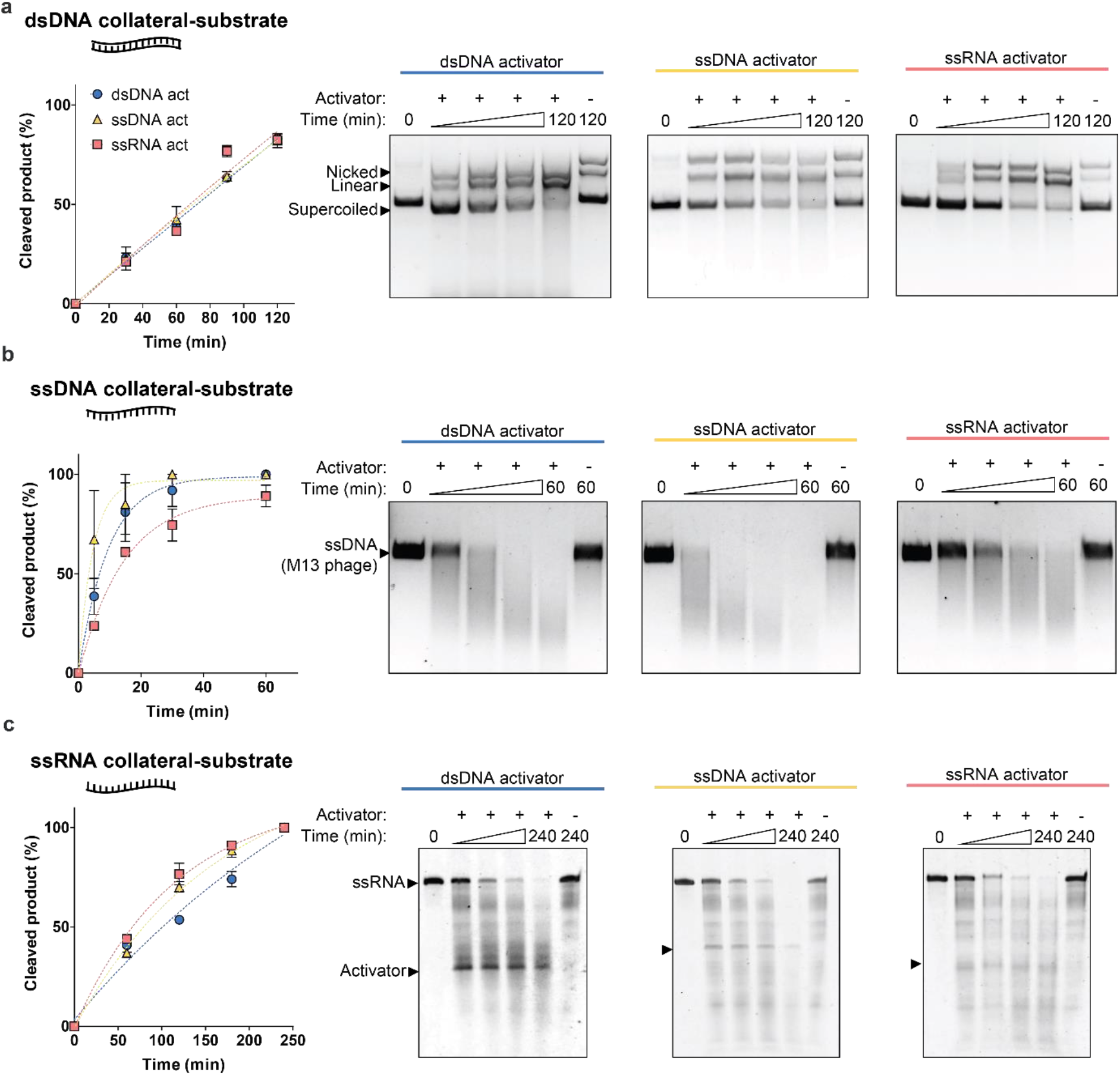
α-synCas exhibits collateral activity against non-target nucleic acids *in vitro*. Time-course of collateral activity against non-target (**a**) dsDNA, (**b**) ssDNA, (**c**) ssRNA by an α-synCas/crRNA complex activated with target dsDNA, ssDNA, and ssRNA substrates. Error bars represent the mean ± SD, where n = 3.

The collateral activity of CRISPR-Cas single effectors has been repurposed for nucleic acid detection in molecular diagnostics^30^. Here, we measured the activity of α-synCas against fluorescent-labelled ssRNA and ssDNA *trans*-substrates as a reporter assay of nucleic acid binding. Preincubation of α-synCas-crRNA binary complex with complementary dsDNA/ssDNA/ssRNA target substrates yielded complexes able to cleave fluorescent-labelled ssRNA (Extended Fig. 8a) and ssDNA (Extended Fig. 8b) probes. The capability of α-synCas to be programmed for detection of specific nucleic acid targets using highly sensitive ssDNA and ssRNA fluorescent probes is unprecedented and opens the possibility of its repurposing for the development of a molecular diagnostic platform capable of recognizing any form of nucleic acid class by using one-single enzyme.

### Structure of α-synCas quaternary complex

With the notable exception of Cas12a2, other natural Cas12 enzymes are unable to cleave dsDNA substrates in *trans*^27,31^. This limitation has been attributed to the RuvC active site not being able to accommodate duplex DNA. In addition, the Nuc domain might also act as a physical barrier limiting cleavage in trans^31^. Analogously to Cas12a2^31^, α-synCas can nick, linearize and degrade supercoiled non-specific plasmid DNA.

The 3.0 Å resolution cryo-EM structure (Fig. 4, PDB: 8QWF; Supplementary Figs. 1 and 2; Extended Table 1) of a quaternary complex of α-synCas bound to crRNA, target dsDNA containing a T-rich PAM, and a non-specific collateral dsDNA, without homology with the crRNA and where both strands contain non-hydrolysable phosphorothioate modifications (Fig. 4a), provides a framework to understand its collateral nuclease activity. While the TS is still hybridized to the crRNA, the cryo-EM density for most of the dissociated NTS is absent, indicating that it has now been displaced from its path in the ternary complex (Fig. 4b and 4c). The net result of this displacement would be to increase the accessibility to the RuvC active site, thus enabling the rapid substrate capture and cleavage in *trans* measured for α-synCas (Fig. 3). The structure captures RuvC in a pre-hydrolysis state during which the non-hydrolysable collateral substrate, visible only at low density thresholds, is attempting to enter its active site (Fig. 4d). This weak density for the collateral substrate is consistent with motion, in agreement with the biochemical data showing dsDNA degradation occurs through multiple-turnover DNA nicking (Fig. 3a).

**Figure 4.**
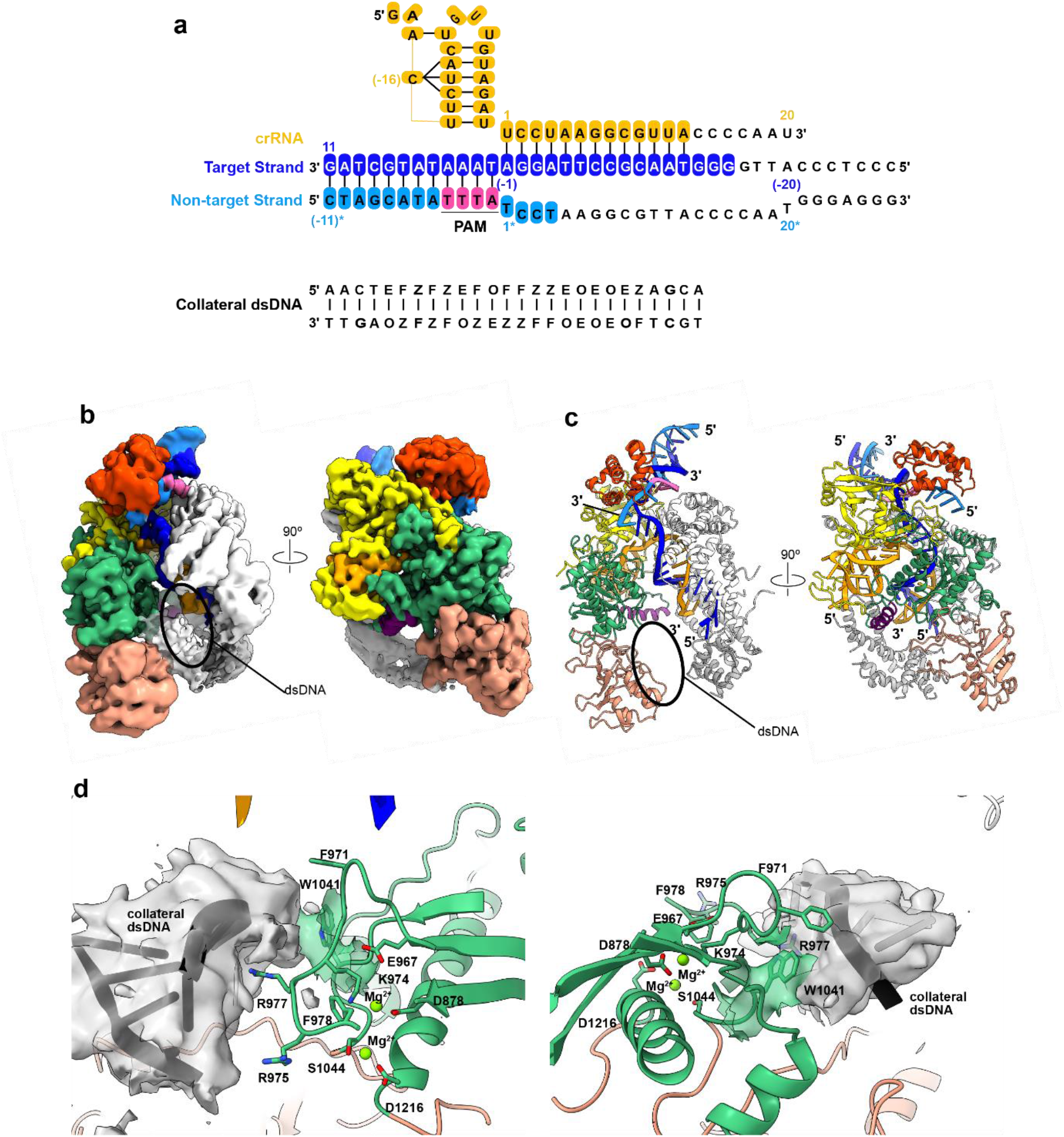
Quaternary complex between α-synCas, crRNA, target dsDNA and collateral dsDNA. **(a)** Sequences of the crRNA guide, target dsDNA and collateral dsDNA used to form the quaternary complex. Nucleotides with coloured background are visible in the cryo-EM map, while uncoloured nucleotides are disordered. The central region of the non-hydrolysable collateral dsDNA includes the following phosphothiates: F, A-Phosphorothioate; O, C-Phosphorothioate; E, G-Phosphorothioate; Z, T-Phosphorothioate. (**b**) Unsharpened cryo-EM map obtained of the quaternary complex coloured by nucleic acid and protein domain. Unassigned density attributed to part of the collateral dsDNA is highlighted. (**c**) Cryo-EM structure of the quaternary complex coloured and oriented as in b. An oval mark the putative location of the collateral dsDNA. (**d**) Close-up views of front (left) and back (right) of the α-synCas active site showing the cryo-EM density attributed to the collateral dsDNA fitted with an idealised four-nucleotide dsDNA fragment. Continuous to the collateral dsDNA density is the density for Trp 1041 shown in transparent green. Selected active site and lid-loop residues are shown as sticks.

### α-synCas exhibits robust editing activity

We demonstrated the ability of α-synCas to efficiently edit DNA in prokaryotes by employing a plasmid interference assay to measure targeting and cleaving performance of the enzyme in *E. coli* cells. This assay measures cell death on selective medium as a result of a resistance-gene encoding plasmid being targeted by the nuclease. Induction of α-synCas/crRNA expression resulted in plasmid depletion and a ∼1000-fold colony reduction in comparison with control conditions without IPTG (Fig. 5a).

**Figure 5.**
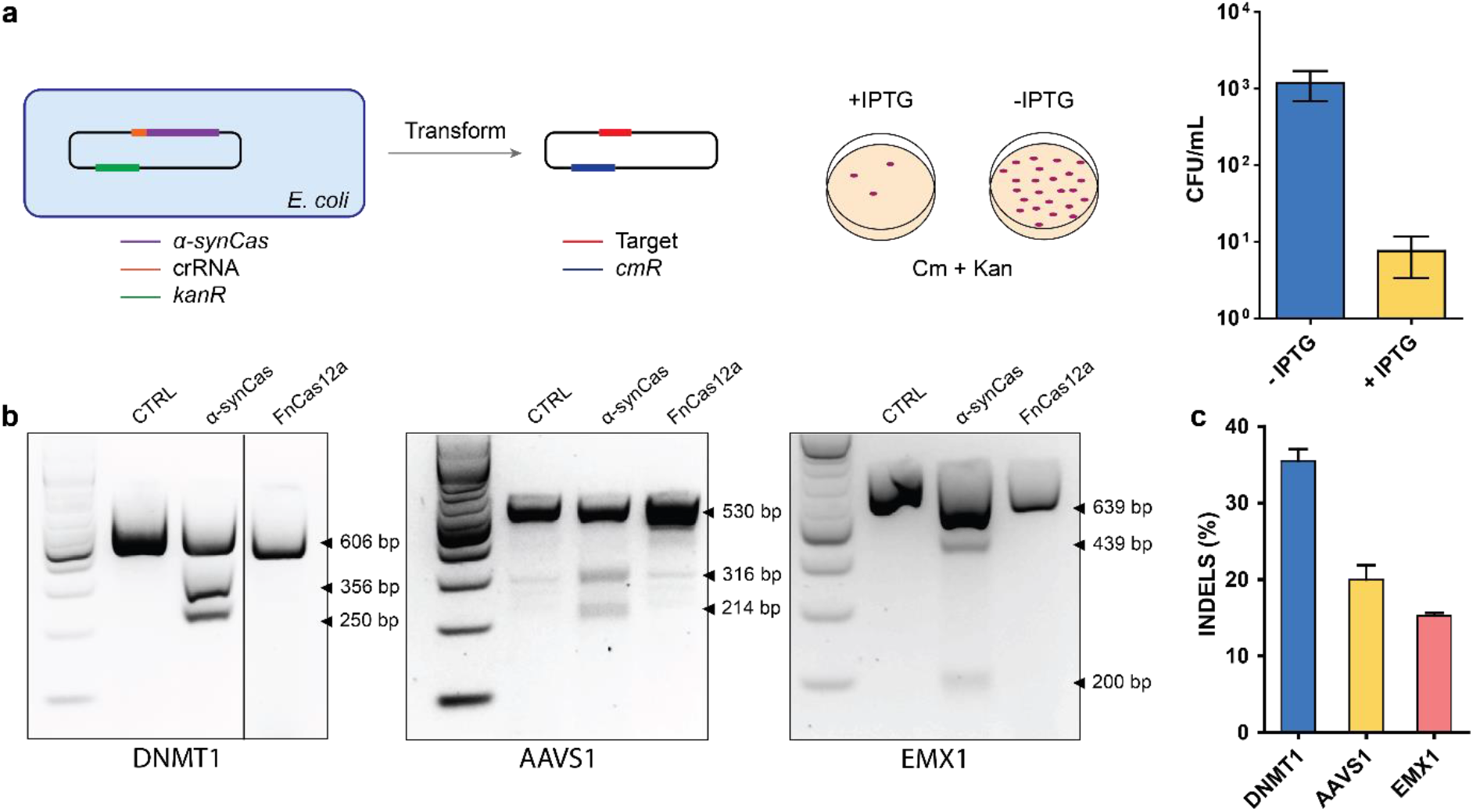
*In cell* editing activity of α-synCas. **(a)** Plasmid clearance assay in *E. coli*. α-synCas expression degrades the target plasmid containing a chloramphenicol resistance gene. Cell survival as a function of α-synCas induction was measured in colony formation units (CFU). (**b**) Editing activity in HEK293T cells. T7 endonuclease mismatch assay for genes *DNMT1*, *AAVS1* and *EMX1* mediated by α-synCas and FnCas12. (**c**) Quantification of indel efficiency achieved by α-synCas at the target sites. Error bars represent the mean ± SD, where n = 3.

Similarly, the genome editing activity of α-synCas in mammalian cells was also robust, as determined by its ability to target, using crRNA guides, the genes *DNMT1, AAVS1* and *EMX1* in HEK293 cells. Compared to FnCas12a, which only exhibited marginal activity against *EMX1* in this assay, α-synCas displayed robust targeting and cleaving activity against all three genes (Fig. 5b). For all the loci tested, the size of the DNA fragments produced as a result of α-synCas action matched those generated by DSB at the expected location. We employed a T7 endonuclease mismatch assay to quantify the occurrence of insertions/deletions (indels), generated by the subsequent cellular non-homologous end-joining repair pathway, with an estimated indel efficiency ranged between 15-35 % for α-synCas (Fig. 5c). Considering that only a limited number of type II and V orthologs have been successfully employed for genome editing in mammalian cells, these results highlight the potential of α-synCas as a gene editing tool.

## Discussion

Protein design techniques using computational methods offer an opportunities to improve and even design new catalysts with properties not found in natural enzymes^32^. The increasing number of protein sequences in databases and the advent of new methods for sequence alterations and design, including deep-learning methods and language models^33–35^, have the potential to expand the universe of biomolecules and revolutionize the fields of biotechnology and synthetic biology. ASR is rapidly gaining prominence in this context, as it provides not only important evolutionary information, but is also able to generate novel synthetic protein sequences not found in nature^36–40^. ASR is arguably the only technique that can handle non-natural sequences with a large number of residues (>1000) and with substantial identity alterations relative to natural proteins. Furthermore, these synthetic proteins provide scaffolds for further engineering and finetuning, thus offering a myriad of possibilities.

We have previously established that ASR can be utilized to uncover important aspects of the evolution of the CRISPR-Cas system^12^. What we present here goes a step beyond, demonstrating that it is possible to obtain a synthetic nuclease with a set of properties that have not been found yet in any existing natural Cas nuclease, surpassing the functionality and versatility of any known CRISPR-Cas effector. As shown before ^12^, ancestral proteins display a functional promiscuity that is clearly present in α-synCas variant, being perhaps its most prominent feature. α-synCas can recognize any nucleic acid form and is able to carry out specific or non-specific *cis/trans* cleavage. This can be done without the need for PAM sequence recognition and with a variety of crRNA-guide sequences. α-synCas can process its own crRNA and is proficient at genome editing in human cells and bacteria. The cryo-EM structures provide a framework to understand the differences between synthetic α-synCas and natural Cas nucleases, and for its finetuning as a biotechnology tool.

In practical terms, the versatility of α-synCas makes it unique and more usable that any other known Cas nuclease, as it can be used for both genome editing and diagnostics based on nucleic acids detection. In recent years, assays based on the collateral activity of Cas12a and Cas13 have been developed as diagnostic tools^30^. Cas12a can only be activated by DNA whereas Cas13 relies on RNA, and thus depending on the sequence to be detected one enzyme or the other must be used. α-synCas is not limited in this way, it can be activated by any nucleic acid and is thus able to identify any type of genetic target. To the best of our knowledge, α-synCas is the only reported nuclease with such expanded activities (Supplementary Table 1). A nuclease with these features might well exist in nature, but searching the natural enzyme space is arduous, costly and time consuming. Similarly, computational methods are still not capable of designing enzymes with the complexity of CRISPR nucleases. These limitations highlight the importance of ASR to generate complex synthetic enzymes with multiple and improved properties.

## Methods

### Engineering of α-synCas sequence

α-synCas was engineered through computational methodologies, specifically, ancestral sequence resurrection (ASR)^12^. We use the BLAST tool with custom parameters and criteria—that is, a maximum of 1,000 hits and minimum identity 35% to ensure the selection of Cas12a sequences (individually inspected) and BLOSUM62 scoring matrix. E-values were virtually zero for all sequences. Sixty-three sequences were selected following similar proportions of sequences in each phylum as in the database (Extended Figure 1). Sequences belong to five bacterial phyla: *Bacillota, Pseudomonadota, Planctomycetota, Spirochaetota* and *Bacteroidota*. Alignment of sequences was performed using MUSCLE software on the MEGA platform and manually edited to eliminate gaps, poorly aligned sites and divergent regions. We inferred the best evolutionary model using MEGA, resulting in the JTT with gamma distribution model (eight categories), Yule model for speciation and length chain of 100 million generations, sampling every 1,000 generations. Phylogeny was carried out using BEAST v.2.6.6 package software (https://beast.community/) including the BEAGLE library for parallel processing and based on Bayesian inference using MCMC. Divergence times were estimated using the Reltime method^41^ implemented in MEGA with discrete eight-category Gamma distribution for evolutionary rates. We set calibration times using information from the TToL^18,42^ in three major clades with 95% confidence interval (CI). Finally, ancestral sequence reconstruction was performed by maximum likelihood using PAML v.4.9 (http://abacus.gene.ucl.ac.uk/software/paml.html) with a gamma distribution of eight categories for variable replacement rates across sites. Posterior probabilities were calculated for all amino acids, and the residue with highest posterior probability was chosen for each site. The reconstructed sequence displays average posterior probability of 0.92 and amino acid sequence identities of 52 % with respect to FnCas12a and 26% with respect to SuCas12a2.

### In vitro characterization

#### Expression and purification of α-synCas

α-synCas gene was synthesized and codon optimized for *E. coli* cell expression. α-synCas was cloned in pET-28a(+) expression vector and transformed in *E. coli* BL21 (DE3) (Life Technologies) for protein expression. Cells were incubated in LB medium at 30 °C at 160 rpm until OD600 reached 0.6 and IPTG (400 µM) was added for protein induction overnight at 18 °C. Cells were pelleted by centrifugation at 5000 g. Pellets were resuspended in extraction buffer (Tris 50 mM, NaCl 500 mM, pH 8, Imidazole 10 mM) supplemented with EDTA Free protease inhibitor (Thermo). Then, the pellet was sonicated for 3 cycles for 10 min at 30% amplitude. Cell debris was separated by ultracentrifugation at 33,000 G for 1 h. For purification, the supernatants were mixed with Ni-NTA agarose beads (Thermo) and incubated for 1 hour. Then, the beads were washed with 50 x column volumes with 40 mM imidazole. After that, imidazole concentration was reduced to 10 mM and aliquot of protease 3C was added. After 1 hour of incubation, protein was eluted in elution buffer (Tris-HCl 50 mM, NaCl 500 mM, pH 8, Imidazole 10 mM). The protein was further purified by size exclusion chromatography using a Superose 6 10/300 GL column (GE Healthcare) and eluted in 50 mM Tris pH 7.5, 150 mM NaCl, 2 mM MgCl_2_. For protein purification verification, sodium dodecyl sulphate– polyacrylamide gel electrophoresis (SDS–PAGE) was used with 8% gels. The protein concentration was calculated by measuring the absorbance at 280 nm in Nanodrop 2000C.

#### crRNA synthesis

crRNA (Supplementary Table 2) was synthesized using a HiScribe T7 High Yield RNA Synthesis Kit (NEB). The DNA sequences includes the T7 promoter at the 5’ end and the sequence from crRNA with the target sequence at the 3’ end. ssDNA oligos were hybridized and the reaction was incubated overnight. Then the crRNA was purified following the protocol of the Monarch RNA Purification Columns Kit.

#### Nucleic acid cleavage assays

For analysis of targeted cleavage (Supplementary Table 3) on supercoiled DNA, 30 µL reactions of 170 nM of α-synCas-crRNA and 100 nM of dsDNA target in 1 x NEB 3.1 buffer (50 mM Tris-HCl pH 7.9, 100 mM NaCl, 10 mM MgCl_2_, 100 µg ml^−1^ BSA) were incubated at 37 °C for varying incubation times. In the case of ssDNA substrate, 30 µL reactions of 80 nM of α-synCas-crRNA and 30 nM of ssDNA target in 1 x NEB 3.1 buffer were incubated at 37 °C for varying incubation times. Reactions were stopped by adding 6X loading dye (NEB) with EDTA and running 1-2% agarose gel. Gels were dyed with SYBR gold (ThermoFisher) and imaged with ChemiDoc XRS + System (Bio-Rad). Cleavage was quantified by ImageJ. Finally, for ssRNA targeted cleavage, 30 µL reactions of 250 nM of α-synCas-crRNA and 120 nM of ssRNA target in 1 x NEB 3.1 buffer were incubated at 37 °C for different varying times. Reactions were stopped by adding 2X loading dye (NEB) with EDTA and running 10% TBE polyacrylamide gels. Gels were dyed with SYBR gold (ThermoFisher) and imaged with ChemiDoc XRS + System (Bio-Rad). Cleavage was quantified by ImageJ.

For analysis of non-targeted (collateral or *trans*) cleavage (Supplementary Table 4, 5), 30 µL reactions of 70 nM of α-synCas-crRNA, 70 nM of target substrate (dsDNA, ssDNA or ssRNA) and 120 nM of collateral substrate (dsDNA, ssDNA or ssRNA) in 1 x NEB 3.1 were incubated at 37 °C for varying incubation times. Reactions were stopped by adding 2X loading dye (NEB) with EDTA or 6X loading dye (NEB) with EDTA and running 10% TBE polyacrylamide gels or 1-2% agarose gel, respectively. Gels were dyed with SYBR gold (ThermoFisher) and imaged with ChemiDoc XRS + System (Bio-Rad). Cleavage was quantified by ImageJ.

#### PAM library construction

A DNA library (Supplementary Table 6, 7) comprising seven random nucleotides was created and subsequently cloned into the plasmid pUC18 by GenScript. This random library was transformed in XL1blue *E. coli* and amplified several times to achieve maximal variability in the PAM sequences. Subsequently, an 855 bp PCR fragment was generated using the primers detailed in Supplementary Table 7 from the DNA library containing the seven random nucleotides.

#### PAM determination

PAM determination assay was performed incubating of 170 nM of α-synCas-crRNA (targeting 23 nt downstream the 7 random nucleotides) and 100 nM of PCR fragment from the DNA library in 1 x NEB 3.1 buffer (50 mM Tris-HCl pH 7.9, 100 mM NaCl, 10 mM MgCl_2_, 100 µg ml^−1^ BSA) for 1 hour at 37 °C. Reaction was stopped added 6X loading dye (NEB) with EDTA and run 2% agarose gel. Gels were dyed with SYBR gold (ThermoFisher) and imaged with ChemiDoc XRS + System (Bio-Rad). The longer fragment, that contained the 7 random nucleotides, was purified from the agarose gel with GeneJet Gel Extraction kit (ThermoFisher) and sequenced by Ion Torrent and the obtained reads were mapped in the reference sequence. Each PAM sequence was quantified, and its frequency calculated from the total PAM previously extracted. From the frequency of each PAM, PAM wheel was generating, following previously published methodology^43^.

For *in vitro* cleavage of different PAM sequences, different DNA fragments carrying each PAM (Supplementary Table 8) were cloned into Zero-Blunt TOPO plasmid. The cleavage assay was performed under the conditions described previously.

#### pre-crRNA processing

Pre-crRNA arrays (Supplementary Table 9) were synthesized using the HiScribe T7 High Yield RNA Synthesis Kit (NEB). T7 transcription was performed for 16 h and then RNA was purified using the Monarch RNA Cleanup Kit (NEB). *In vitro* cleavage was performed with purified recombinant. α-synCas (330 mM) and in vitro-transcribed pre-crRNA arrays (100 nM) were incubated at 37 °C 1 x NEB 3.1 buffer for 1 hour. Reactions were stopped by adding 2X loading dye (NEB) with EDTA and running 10% TBE polyacrylamide gels. Gels were dyed with SYBR gold (ThermoFisher) and imaged with ChemiDoc XRS + System (Bio-Rad).

### Nucleic acid detection by α-synCas with ssRNA and ssDNA reporter probes

For nucleic acid detection with fluorescent reporter probes, 70 nM of α-synCas-crRNA complex was combined with RNase or DNase Alert (250 nM, IDT) and 70 nM of target substrate (dsDNA, ssDNA or ssRNA) in a 96-well plate (Corning). The reactions were incubated at 37 °C and monitored for reporter fluorescence (RNase Alert: excitation 490 / emission 520, DNAse Alert: excitation 530/emission 560) using the multi-mode microplate reader (BioTek Instruments). A negative control was prepared with nuclease-free water instead of nucleic acid target.

### In cell characterization

#### Plasmid clearance assay in E. coli

Plasmid clearance assay was performed in *E. coli* BL21 containing nuclease (α-synCas) and crRNA-expressing plasmid (pET-28a(+)). Bacterial culture was grown overnight and used to inoculate fresh ZymoBroth with 50 μg/mL kanamycin to an O.D. of 0.4–0.6 and made competent using a Mix & Go kit (Zymo). Immediately after, the competent *E. coli* pool was transformed with 500 ng of the target plasmid (pAF plasmid: a modified version of pBAD33 plasmid containing a subcloned LacZ gene, Supplementary Table 10) and incubated on ice for 15 min. The cells were recovered for 2 h at 37 °C with shaking in 500 µL of LB without antibiotics. Next, the cultures were sequentially diluted, and the dilutions were plated on LB agar plates containing 50 μg/mL kanamycin and 25 μg/mL chloramphenicol in the absence or presence of IPTG (0.3 mM).

#### Endogenous gene editing in human HEK293T cells

Human HEK293T cells (ATCC) were maintained in DMEM (Gibco) supplemented with 10% heat inactivated FBS (Gibco) in an incubator at 37 °C and 5% CO2. HEK293T cells were seeded onto 24-well plates (Thermo) 24 h before transfection. Cells were transfected using jetPrime (Polyplus) at 70–80% confluency following the manufacturer’s recommended protocol. For each well of a 6-well plate, a total of 500 ng plasmid DNA (humanized version of α-synCas cloned into pcDNA3.1) and 500 ng of PCR amplicons comprised of a U6 promoter driving expression of the crRNA for targeted each gene were used (Supplementary Table 11).

HEK293T cells were transfected with DNA, as described above. Cells were incubated at 37 °C for 48 h after transfection before genomic DNA extraction. Genomic DNA was extracted using the GENELUTE™ mammalian genomic DNA Miniprep kit (Merck) following the manufacturer’s protocol. Editing activity of α-synCas was evaluated using EnGen Mutation Detection Kit (NEB). Briefly, the genomic region flanking the target site for each gene was PCR amplified (Supplementary Table 11) using Q5® Hot Start High Fidelity and PCR products were subjected to a re-annealing process to enable heteroduplex formation. After re-annealing, products were treated with T7 endonuclease l, and analyzed on 2 % agarose gels. Gels were dyed with SYBR gold (ThermoFisher) and imaged with ChemiDoc XRS + System (Bio-Rad). Cleavage was quantified by ImageJ. Quantification was based on relative band intensities. Indel percentage was determined by the formula,

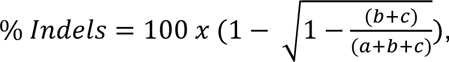

where *a* is the integrated intensity of the undigested PCR product band, and *b* and *c* are the integrated intensities of each cleavage product band.

### Cryo-EM structural determination

#### Cryo-EM data collection

For all specimens (see Supplementary Note 2 for details on specimen preparation), immediately after adding 0.05% CHAPS, 4 ul aliquot of the mixture was adsorbed onto a glow-discharged Quantifoil R 1.2/1.3 300 mesh grid (Quantifoil) and vitrified in liquid ethane with a Leica EM GP2 cryoplunger (Leica) using front-side blotting for 2 s at 95% humidity. The vitrified complex specimens were imaged in house using a 300 kV Krios G4 (ThermoScientific) equipped with a BioContinuum/K3 camera (Gatan) operating in counting mode at a calibrated 0.8238 Å/pix. Employing a 1-1.6 μm underfocus range we recorded three movies per hole with a total accumulated dose of 50 e^−^/Å^2^ over 50 frames. Movies were recorded automatically using EPU 2 (ThermoScientific) with Aberration-free image shift (AFIS) and Fringe-free imaging (FFI).

#### Cryo-EM data processing

Initial processing of complex specimen data followed the same general processing strategy (Supplementary Figs. 1 and 3). Initial frame alignment and contrast transfer function (CTF) estimation was performed using cryoSPARC live^44^. Following processing steps were carried out in cryoSPARC^44^. (See Supplementary Figs. 1-3 and Supplementary Note 2 for details on the cryo-EM data processing and 3D reconstruction).

#### Model building

Initial models were obtained using automated model building with ModelAngelo^45^, and completed by manual intervention in Coot^46^ (See Supplementary Note 2 for details). To improve the fit and to optimize stereochemistry of the atomic model real space refinements with secondary structure and geometry restraints were performed using Phenix^47^.

### Statistics and reproducibility

Results are representative of least three independent experiments. All replication attempts showed similar results.

## Data Availability

We have made available sequencing data. Other data supporting the findings of this study are available from the corresponding authors upon reasonable request. Sequencing data for PAM determination and gene editing experiments will be made available through the National Center for Biotechnology Information Sequence Read Archive (NCBI SRA) under BioProject ID XXXX. The EM maps of apo α-synCas, accession numbers EMD-18691 and EMD-18692, and its ternary and quaternary complexes, EMD-18693 and EMD-18694 have been deposited in the Electron Microscopy Data Bank (http://www.ebi.ac.uk/pdbe/emdb/). The atomic coordinates of α-synCas, and its ternary and quaternary complexes, have been deposited in the Protein Data Bank (www.pdb.org) under PDB ID codes 8QWD, 8QWE and 8QWF, respectively.

## Acknowledgments

This work has been supported by grants PID2019-109087RB-I00 and PID2022-141347OB-I00 to R.P.-J, PID2019-104423GB-I00 and PID2022-143177NB-I00 to I. U.-B, PID2021-126263OA-I00 to G. A-P. from Spanish Ministry of Science and Innovation. This project has received funding from the European Union’s Horizon 2020 research and innovation programme under grant agreement No 964764 to R.P.-J. The content presented in this document represents the views of the authors, and the European Commission has no liability in respect to the content. Y.J. acknowledges financial support from Juan de la Cierva program grant FJC2021-047689-I from Spanish Ministry of Science and Innovation. R.P.-J. and Y.J. acknowledge financial support from Spanish Foundation for the promotion of research of Amyotrophic Lateral Sclerosis (FUNDELA). CIC bioGUNE support was provided from The Department of Industry, Tourism and Trade of the Government of the Autonomous Community of the Basque Country (Elkartek Research Programs 2020-2023), the Innovation Technology Department of the Bizkaia County and MINECO for the Severo Ochoa Excellence Accreditation (CEX2021-001136-S). CIBERehd is funded by the Instituto de Salud Carlos III. High-resolution cryo-EM data collection was performed at the Basque Resource for Electron Microscopy supported primarily by the Department of Education and the Innovation Fund of the Basque Government, the Fundación Biofísica Bizkaia, and the Spanish Ministry of Science and Innovation, through the Plan de Recuperación, Transformación y Resiliencia (PRTR) funded by the NextGenerationEU (PRTR-C17.I1) program. We acknowledge to Prof. Matxalen Llosa-Blas for kindly providing the target plasmid for plasmid clearance assay.

## Author contributions

R. P-J. conceived the project. R. P-J. and Y. J. designed research and planned experiments. Y. J. and R. P-J. performed the phylogenetic analysis and ancestral sequence reconstruction. Y. J, and S.S., cloned and expressed proteins and performed in vitro experiments, functional validation of α-synCas in mammalian and bacterial cells. Y. J., M.G-L and A.M.A. performed the sequencing experiments and bioinformatic analysis. I. T, J.P. L-A., G. A-P. and I. U-B. designed, performed, analysed and represented structural data. All authors participated in discussions and provided ideas for the work. Y. J., I.T., I. U.-B. and R.P-J. wrote the manuscript with additional input end editions from all authors.

## Additional information

**Extended data** is available for this paper at…

**Supplementary information** The online version contains supplementary material available at..

**Correspondence and requests for materials should be addressed** to Iban Ubarretxena-Belandia or Raul Perez-Jimenez

---

**Extended Figure 1.**
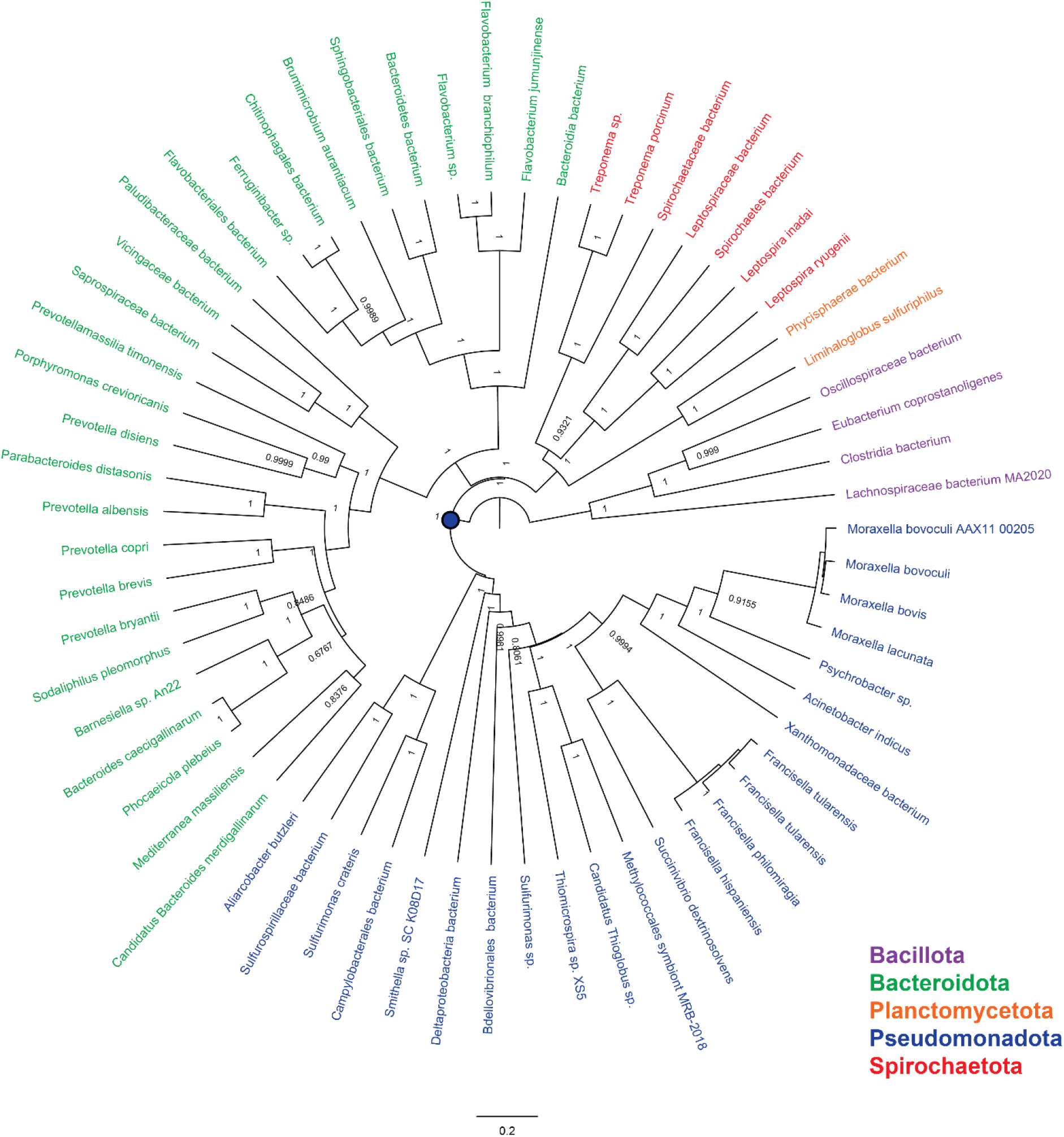
Phylogenetic tree of sequences used for ASR of α-synCas. Posterior probabilities for each node are indicated. A blue circle marks the node selected for ancestral sequence reconstruction.

**Extended Figure 2.**
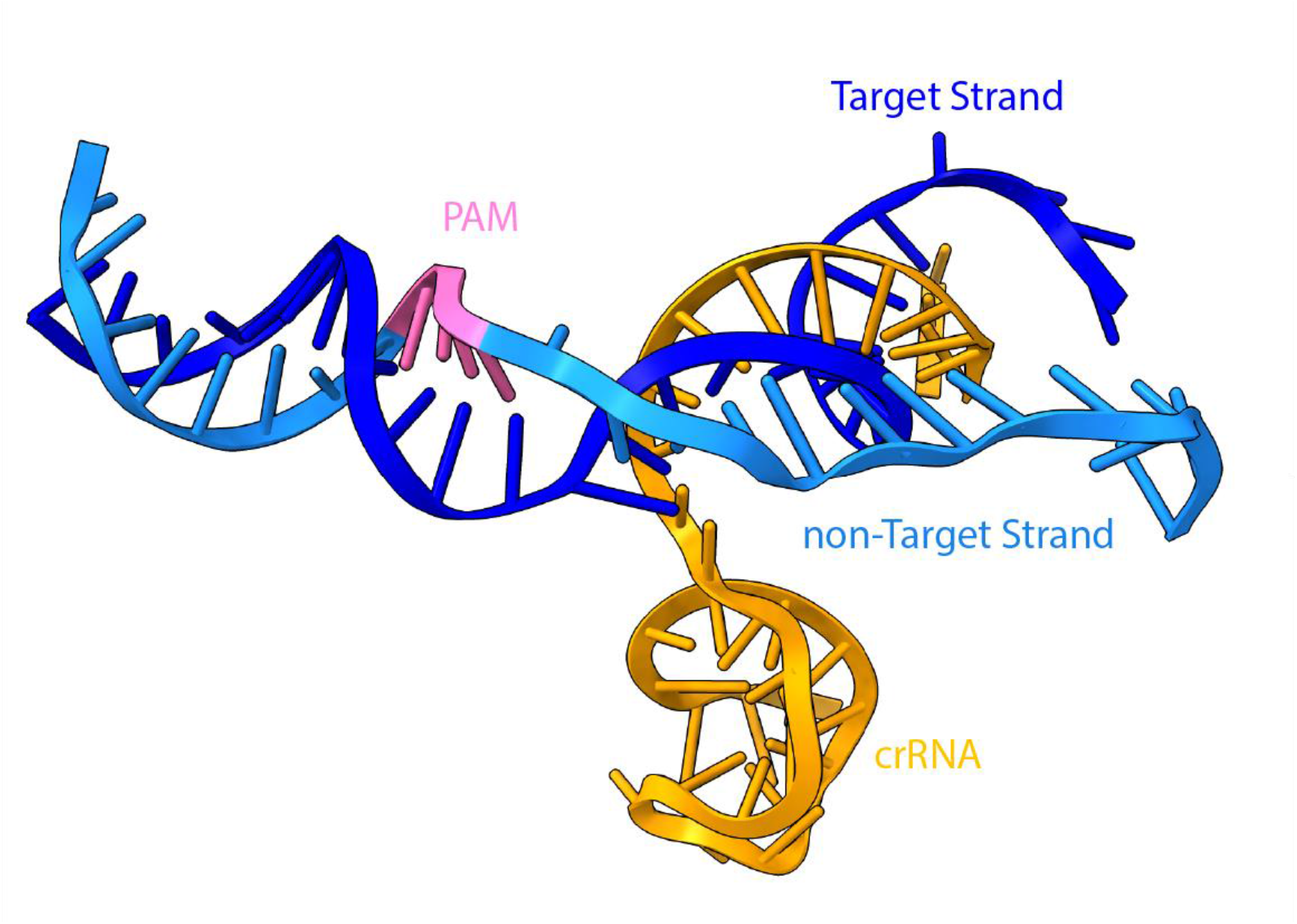
Structures of the crRNA and target dsDNA in the ternary complex.

**Extended Figure 3.**
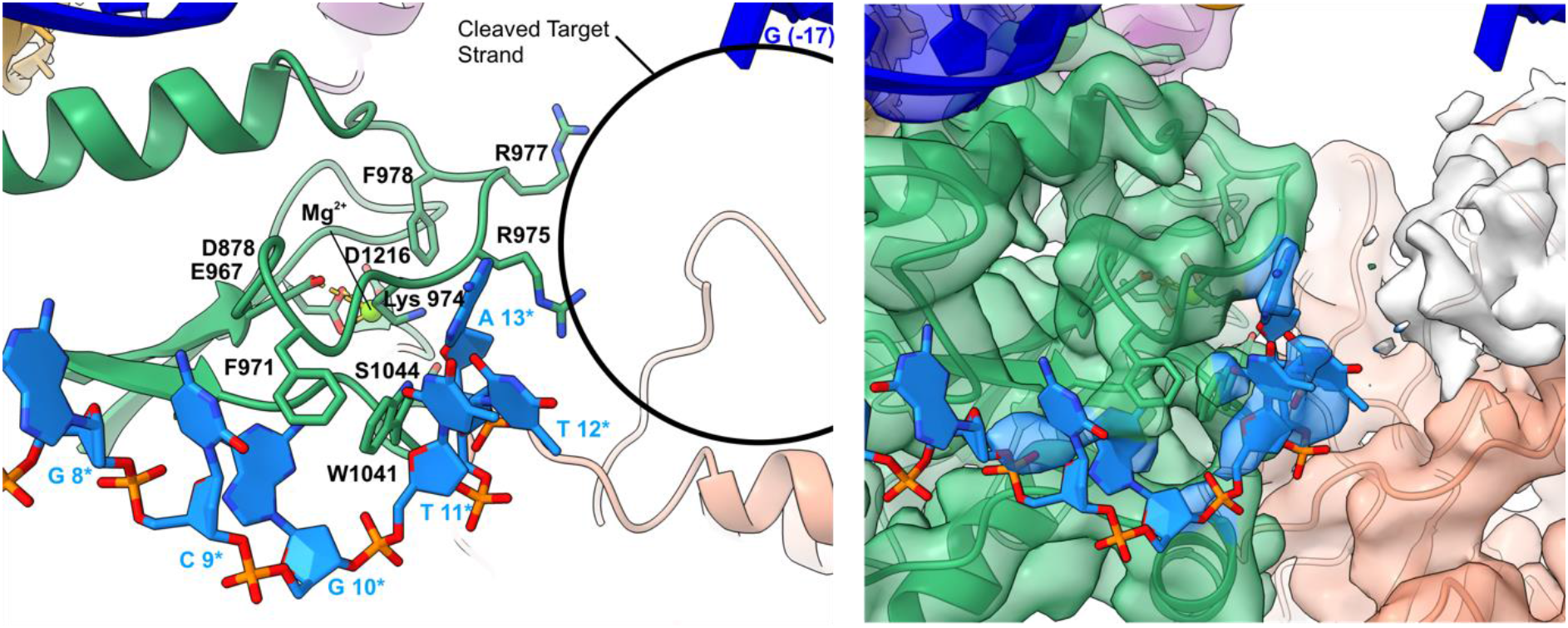
Close-up of the ternary complex active site. On the left, the active site and lid-loop residues are shown as sticks and the non-target DNA strand nucleotides as filled sticks. A circle marks the putative location of the cleaved target strand DNA. On the right, the fitting of the active site region on the cryo-EM density map.

**Extended Figure 4.**
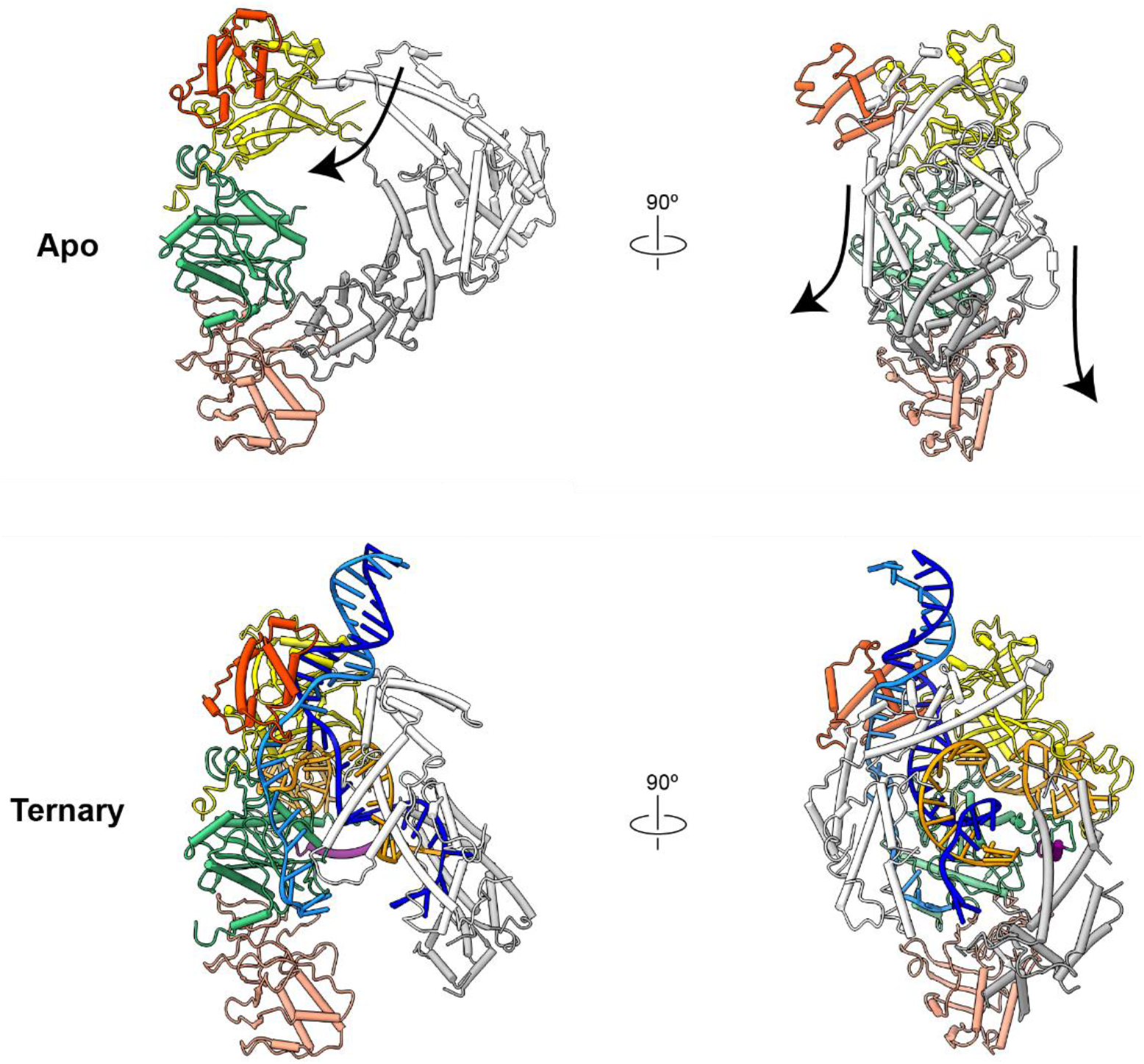
Cryo-EM structure of apo α-synCas. Comparison of apo (top) and ternary complex (bottom) structures of α-synCas coloured by nucleic acid and protein domain. Arrows depict the direction of movement of the REC1 and REC2 domains upon RNA/DNA binding.

**Extended Figure 5.**
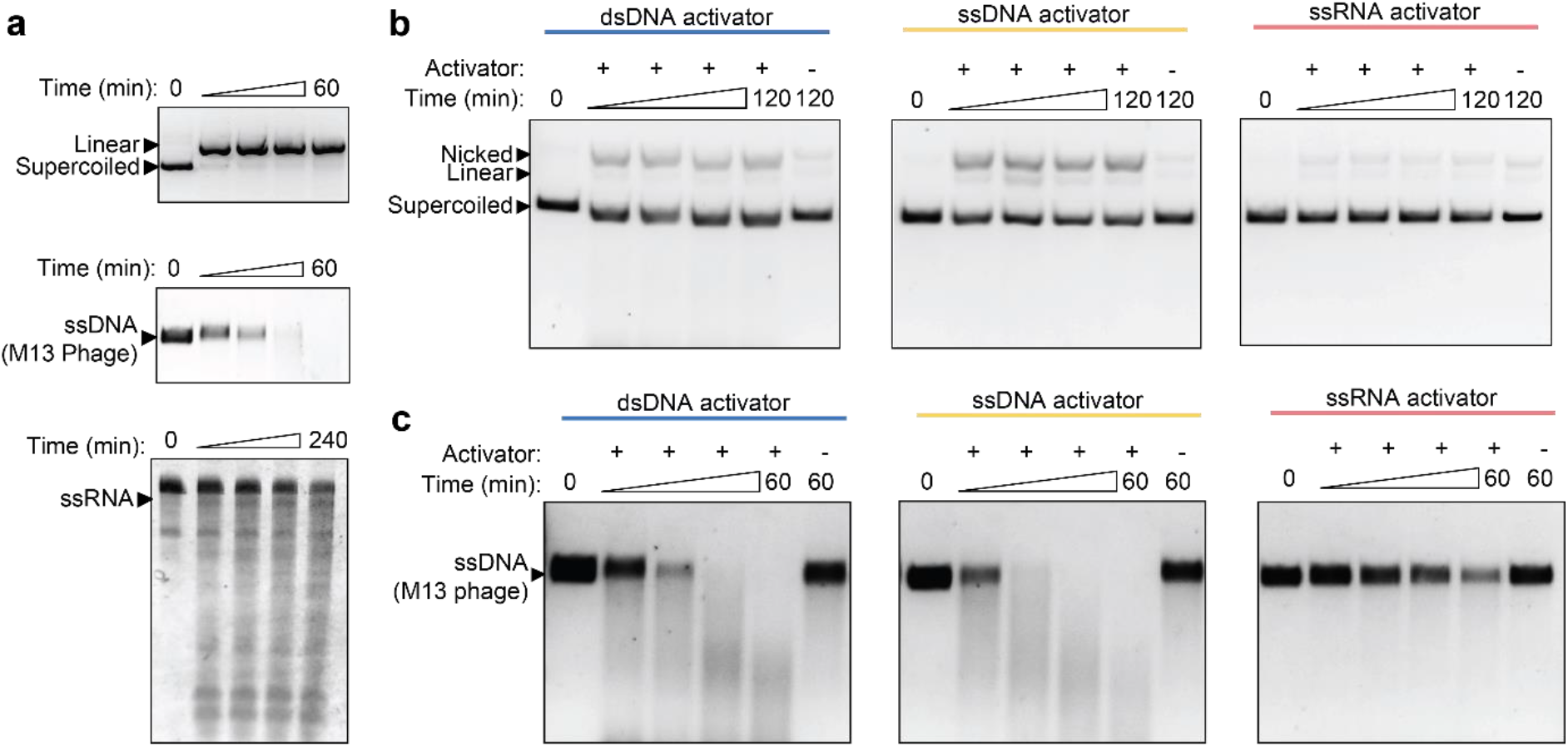
Activities of FnCas12a ortholog. (**a**) Cleavage activity of FnCas12a against target dsDNA, ssDNA and ssRNA. (**b**) Collateral cleavage activity against non-target dsDNA and (**c**) ssDNA by an FnCas12a/crRNA complex activated by target nucleic-acid substrates.

**Extended Figure 6.**
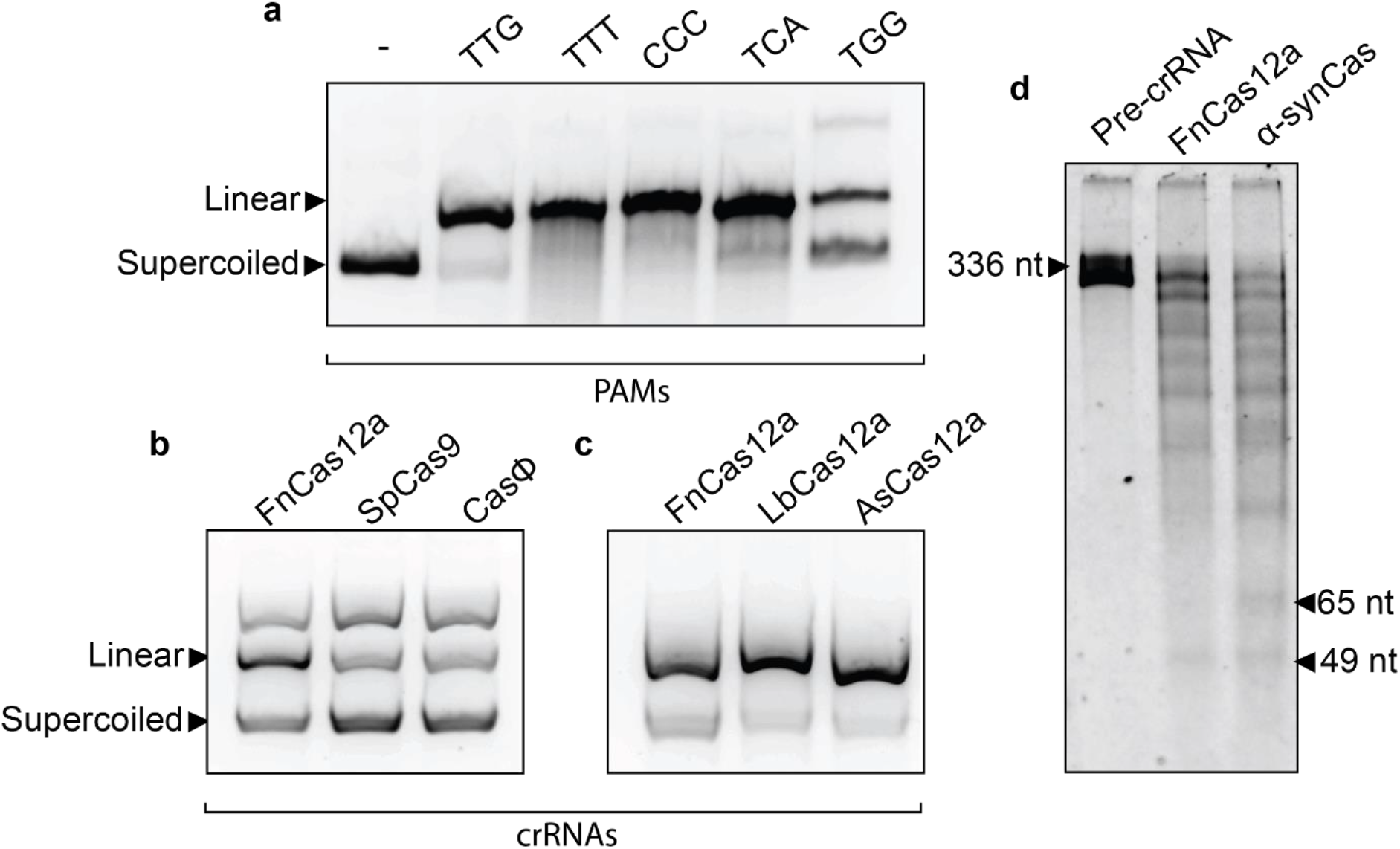
Promiscuous activities of α-synCas. (a) In vitro activity against supercoiled DNA as a function of PAM sequence variation. (**b**) *In vitro* cleavage assay against supercoiled DNA as a function of crRNAs from different CRISPR types and species (**c**). (**d**) *In vitro* processing of FnCpf1 pre-crRNA transcript with purified FnCas12 and α-synCas.

**Extended Figure 7.**
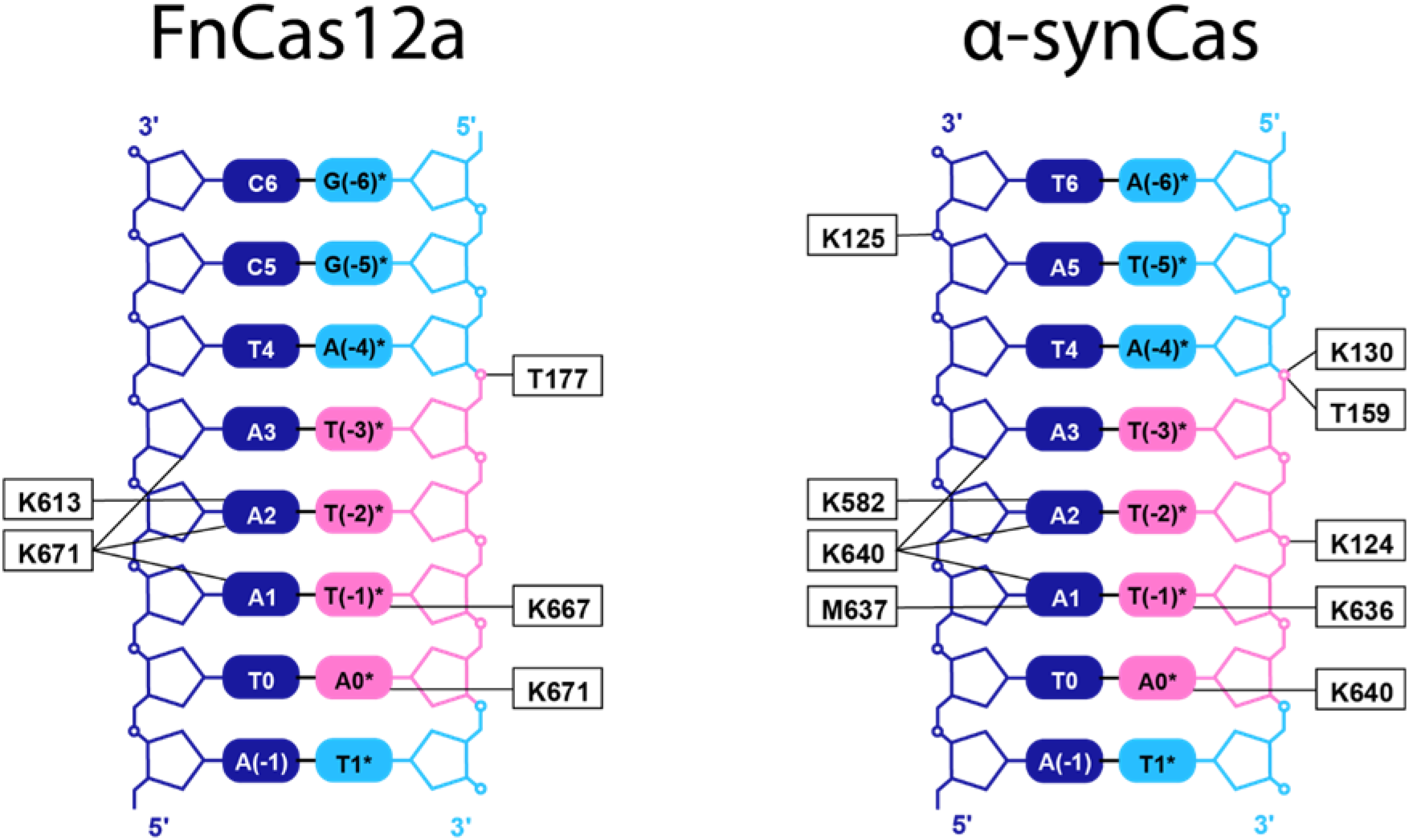
Schematic of the interactions of target and non-target DNA strands with FnCas12a (left) and α-synCas (right). Nucleotides are coloured as in Figure 1b.

**Extended Figure 8.**
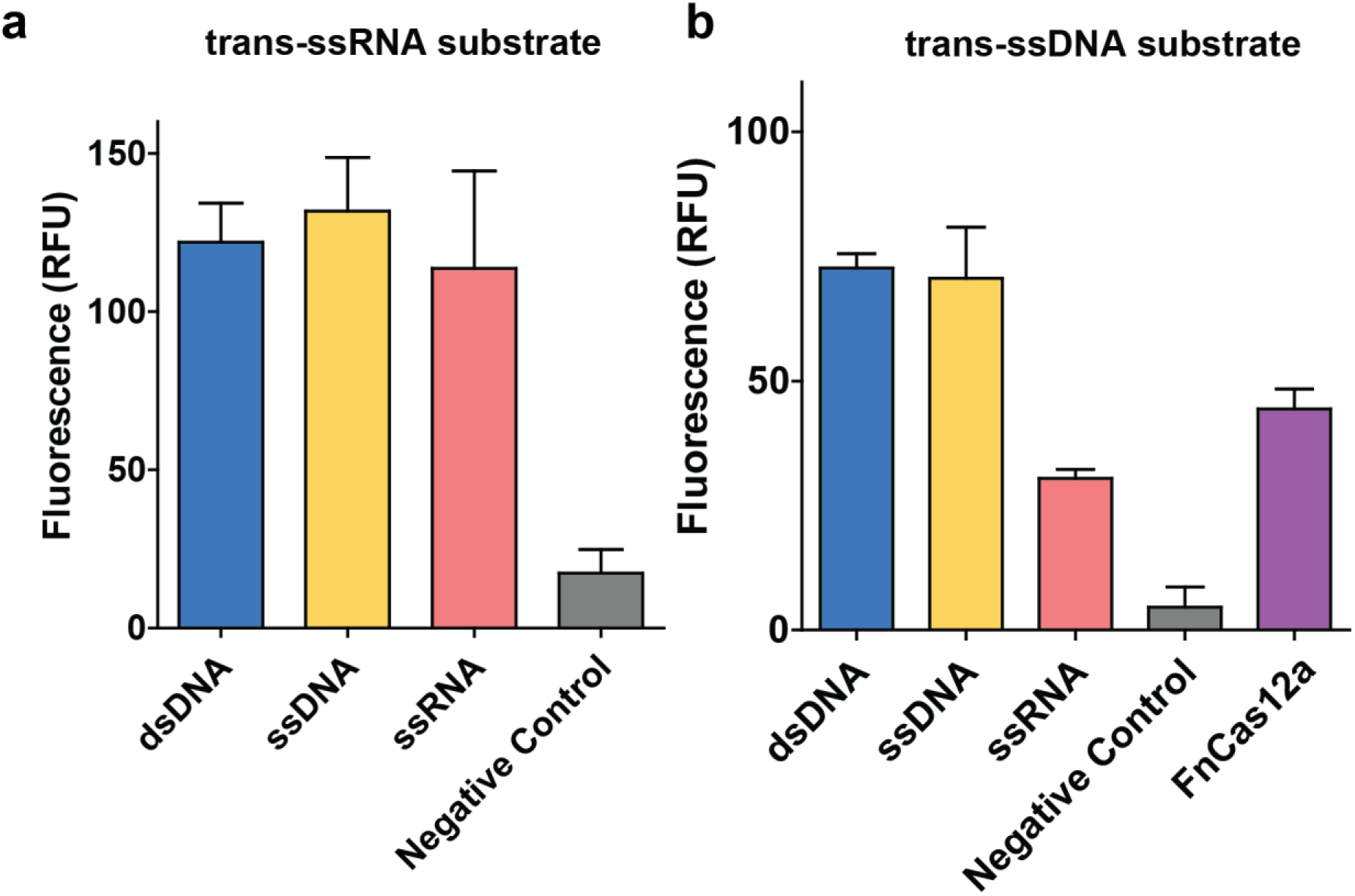
Nucleic acid detection assays. Quantification of maximum fluorescence signal generated after incubating α-synCas-crRNA-activator with a custom (**a**) *trans*-ssRNA-FQ or (**b**) *trans*-ssDNA-FQ reporter. Error bars represent the mean ± SD, where n = 3.

**Extended Table 1.** Cryo-EM data collection, refinement and validation statistics.

| | Apo α-synCas<br>(EMD-18691)<br>(PDB 8QWD) | Ternary complex<br>α-synCas-crRNA-target dsDNA<br>(EMD-18693)<br>(PDB 8QWE) | Quaternary complex<br>α-synCas-crRNA-target dsDNA-collateral dsDNA<br>(EMD-18694)<br>(PDB 8QWF) |
| --- | --- | --- | --- |
| **Data collection and processing** |  |  |  |
| Magnification | 105000 | 105000 | 105000 |
| Voltage (kV) | 300 | 300 | 300 |
| Electron exposure (e-/Å<sup>2</sup>) | 50.979 | 50.979 | 50.084 |
| Defocus range (μm) | 1-1.6 | 1-1.6 | 1-1.6 |
| Pixel size (Å) | 0.8238 | 0.8238 | 0.8238 |
| Symmetry imposed | C1 | C1 | C1 |
| Initial particle images (no.) | 3324256 | 5463714 | 7589433 |
| Final particle images (no.) | 254316 | 137913 | 300509 |
| Map resolution (Å) | 3.33 | 3.14 | 3.08 |
| FSC threshold | 0.143 | 0.143 | 0.143 |
| Map resolution range (Å) | 2.9-6.5 | 2.9-6.5 | 2.9-6.5 |
| **Refinement** |  |  |  |
| Initial model used (PDB code) | - | - | - |
| Model resolution (Å) | 3.3 | 3.1 | 3.0 |
| FSC threshold | 0.143 | 0.143 | 0.143 |
| Model resolution range (Å) | 2.9-6.5 | 2.9-6.5 | 2.9-6.5 |
| Map sharpening *B* factor (Å<sup>2</sup>) | -137.6 | -103.2 | -124.0 |
| Model composition |  |  |  |
| Non-hydrogen atoms | 9204 | 11713 | 11499 |
| Protein residues | 1103 | 1185 | 1186 |
| Ligands | 2 | 5 | 6 |
| *B* factors (Å<sup>2</sup>) |  |  |  |
| Protein | 127.16 | 101.79 | 31.38 |
| Nucleotide | - | 93.27 | 35.04 |
| Ligand | 69.59 | 28.99 | 21.31 |
| R.m.s. deviations |  |  |  |
| Bond lengths (Å) | 0.002 | 0.005 | 0.002 |
| Bond angles (°) | 0.507 | 0.645 | 0.499 |
| Validation |  |  |  |
| MolProbity score | 1.79 | 1.63 | 1.69 |
| Clashscore | 8.29 | 6.94 | 7.59 |
| Poor rotamers (%) | 0.20 | 0.00 | 0.00 |
| Ramachandran plot |  |  |  |
| Favored (%) | 95.02 | 96.33 | 95.99 |
| Allowed (%) | 4.98 | 3.67 | 4.01 |
| Disallowed (%) | 0.00 | 0.00 | 0.00 |

