## Supplementary material for "A synthetic CRISPR-Cas nuclease with expanded enzymatic activities": Supp Information

**Table of Contents**

**Supplementary Figures**

Supplementary Figure 1. Cryo-EM data processing pipeline for ternary and quaternary complex.
Supplementary Figure 2. Local resolution of cryo-EM reconstructions.
Supplementary Figure 3. Cryo-EM data processing pipeline for apo-α-synCas.
Supplementary Figure 4. Blocked active site of apo α-synCas.

**Supplementary Tables**

Supplementary Table 1. Features of the main types of Class II CRISPR-Cas effectors compared to α-synCas.
Supplementary Table 2. crRNA sequences for in vitro experiments.
Supplementary Table 3. Target sequences for targeted cleavage experiments.
Supplementary Table 4. Activators sequences (CIS-substrates) for non-targeted cleavage experiments.
Supplementary Table 5. Trans substrates for non-targeted cleavage experiments.
Supplementary Table 6. PAM library cloned in pUC18. Target sequence is indicated in blue and random PAM in green.
Supplementary Table 7. Primers for PAM determination assay.
Supplementary Table 8. In vitro cleavage PAM sequences. Target sequence indicated in blue and PAM sequence in green.
Supplementary Table 9. Sequences of pre-crRNA transcript.
Supplementary Table 10. Sequences for plasmid clearance assay in E. coli.
Supplementary Table 11. Loci sequences and primers used for endogenous gene editing in human HEK293T cells.
Supplementary Table 12. crRNAs and oligos used for cryo-EM analysis.

**Supplementary Notes**

Supplementary Note 1. Structure of apo α-synCas.
Supplementary Note 2. Detailed Cryo-EM materials and methods.

**References**

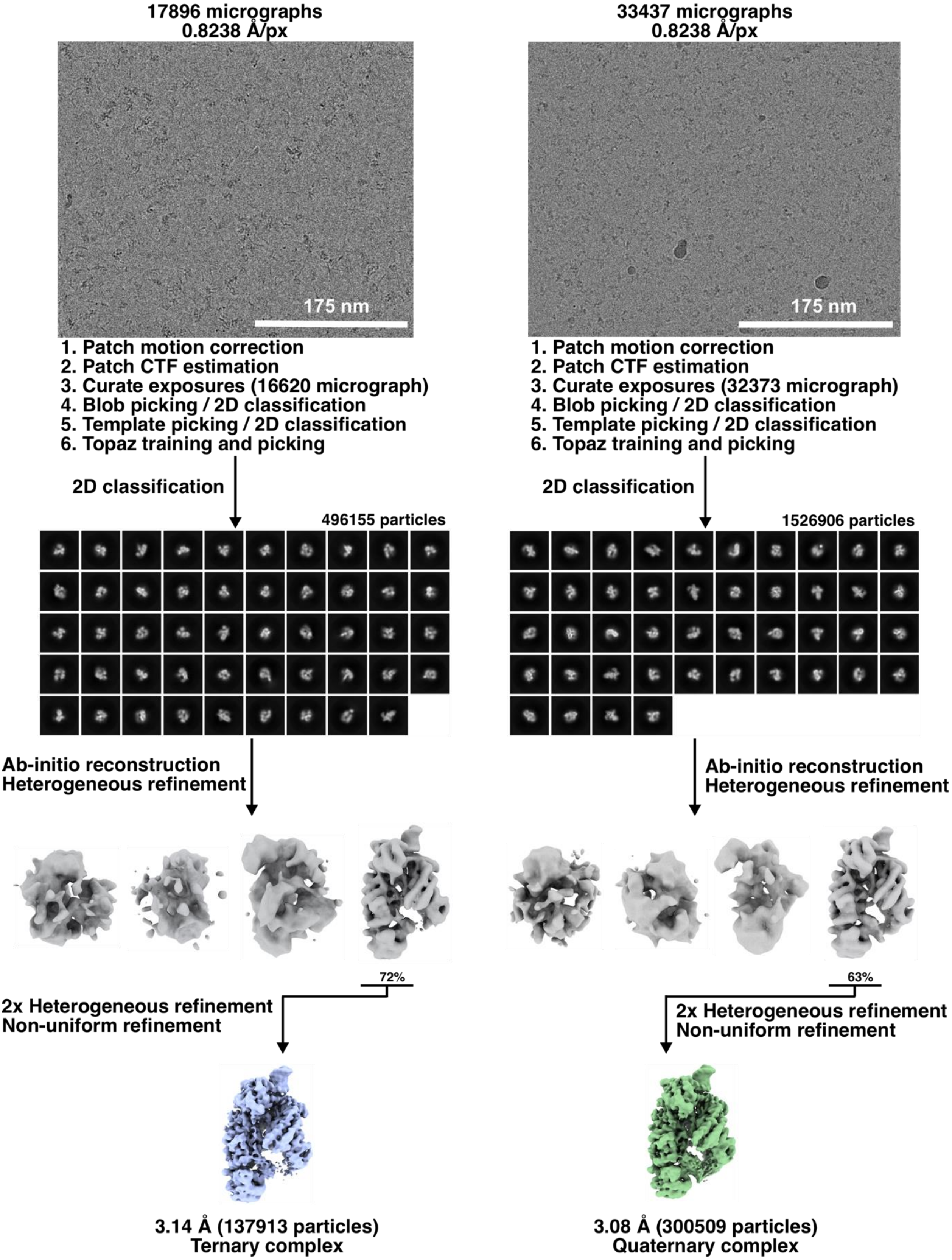


**Supplementary Figure 1. Cryo-EM data processing pipeline for ternary and quaternary complex.**

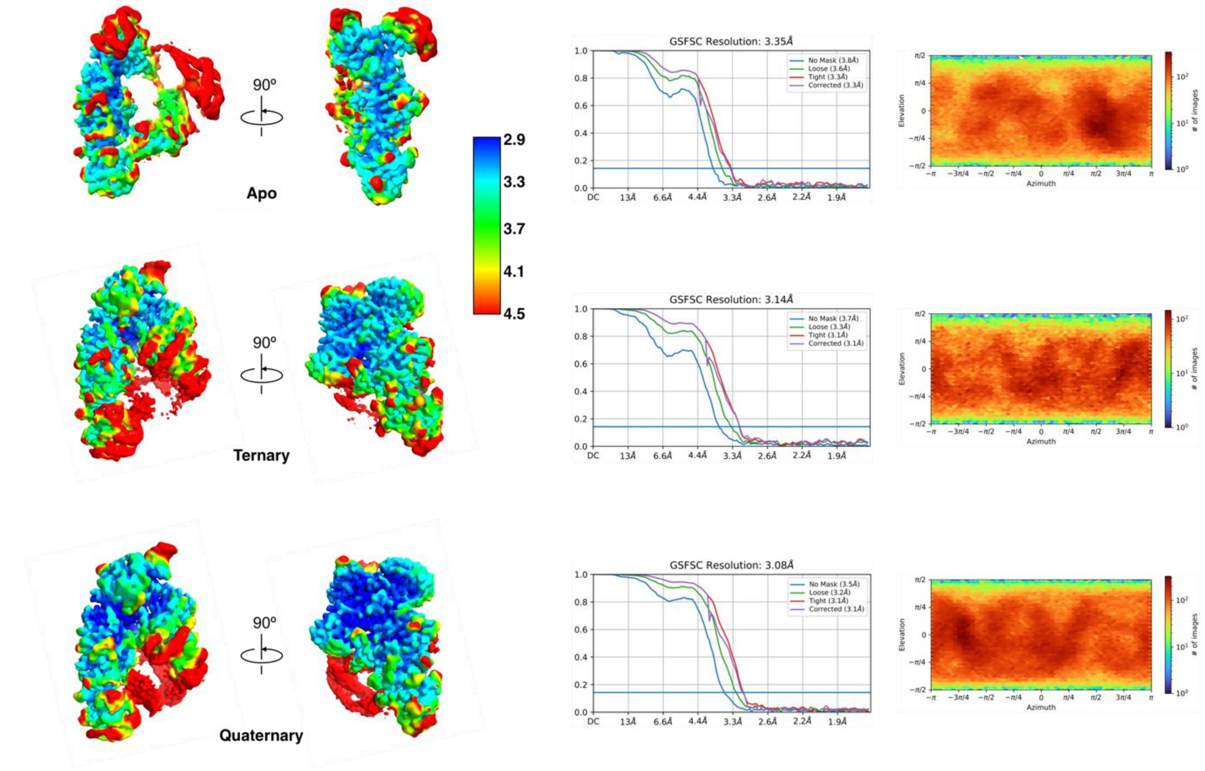


**Supplementary Figure 2. Local resolution of cryo-EM reconstructions.** Unsharpened maps of $\alpha$-synCas complexes colored by local resolution (left). Gold standard FSC curves for each reconstruction. Estimated resolution at FSC = 0.143 (middle). Euler diagrams for each reconstruction showing particle orientation distribution (right).

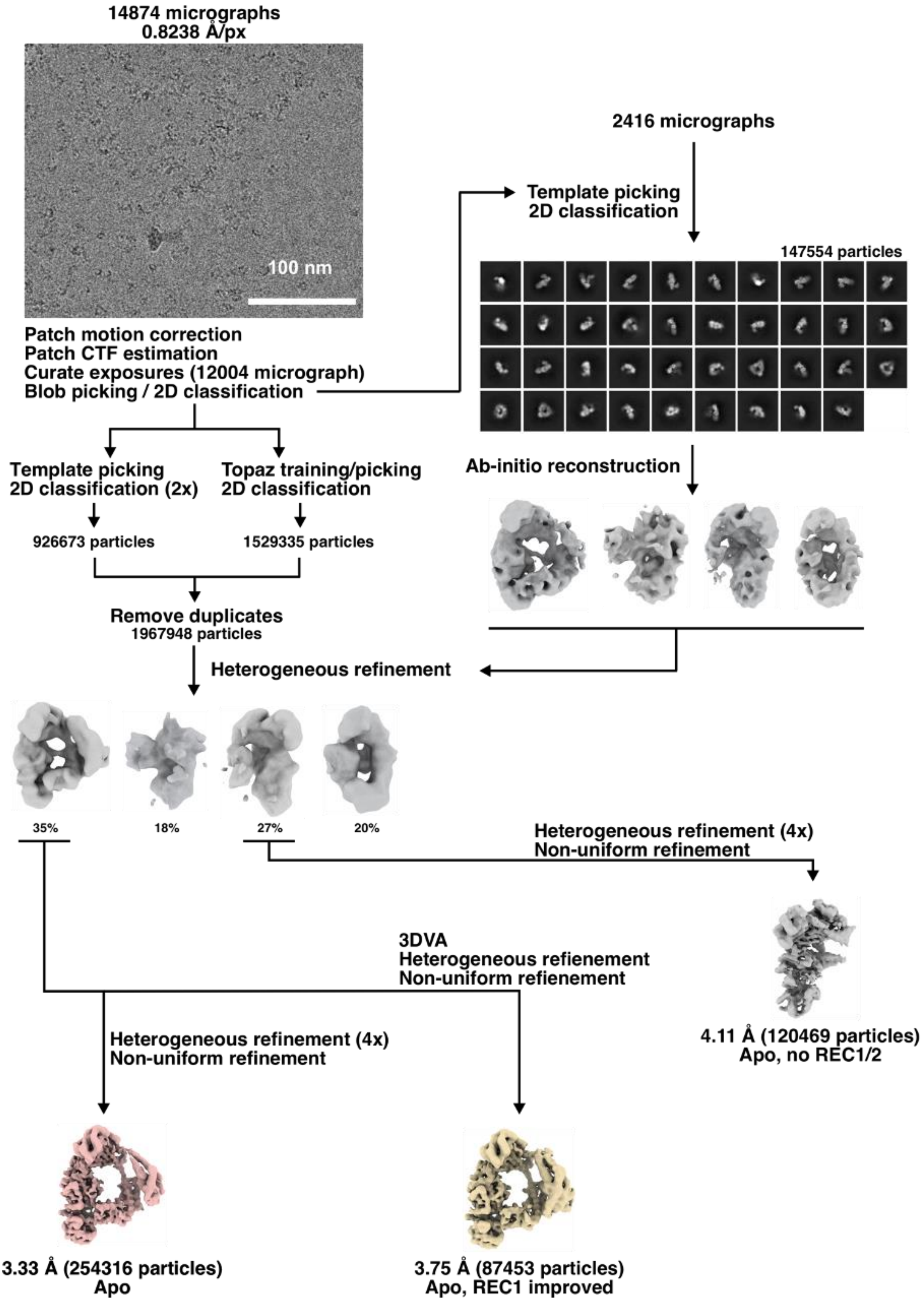


**Supplementary Figure 3. Cryo-EM data processing pipeline for apo-$\alpha$-synCas.**

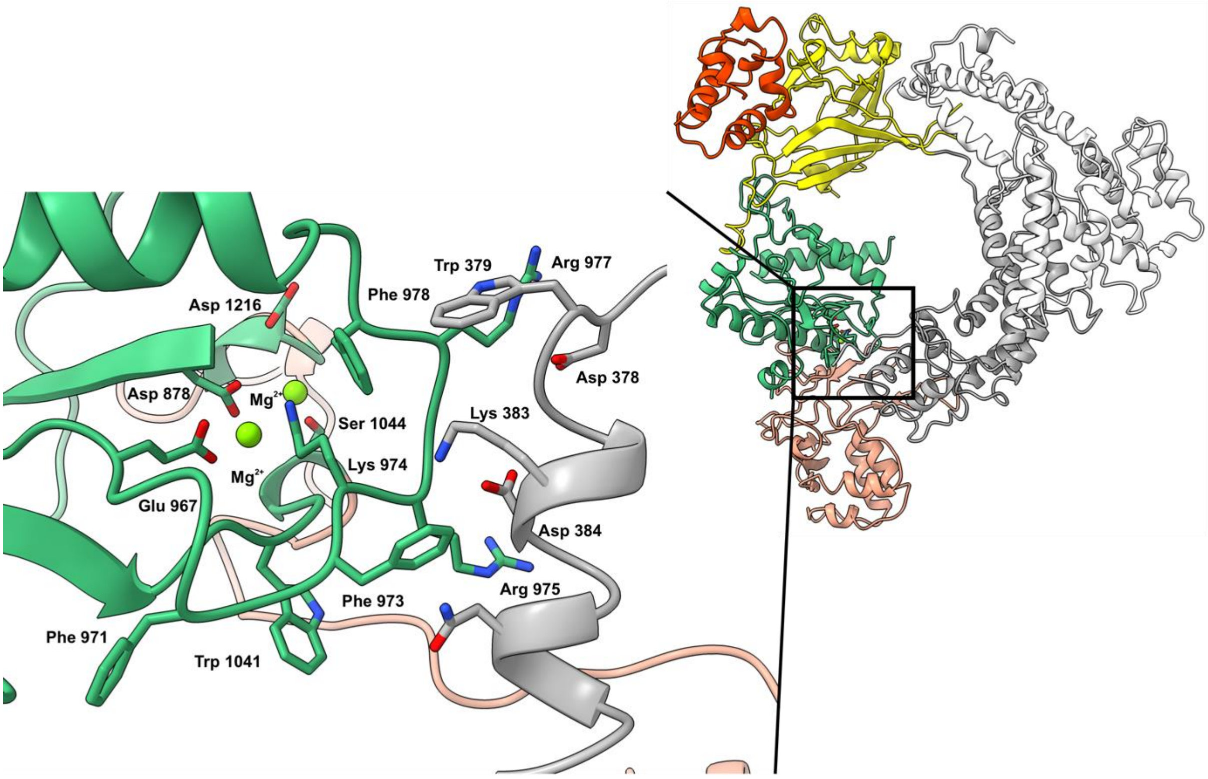


**Supplementary Figure 4**. **Blocked active site of apo α-synCas.** Overall structure of the apo state colored by protein domain as in **Fig 1a**. The inset panel displays a close-up view to the α-synCas active site, which is blocked by REC2. Interacting residues of the REC2 domain (in grey), and residues of the RuvC domain (in green) are shown as sticks. Residues in the active site are shown as sticks and the $Mg^{2+}$ ions as green spheres.

**Supplementary Table 1. Features of the main types of Class 2 CRISPR-Cas effectors compared to α-synCas.**

| Characteristics | Cas9 | Cas12a | Cas13 | α-synCas |
|---|---|---|---|---|
| **Pre-crRNA processing** | No | Yes | No | **Yes** |
| **tracrRNA** | Yes | No | No | No |
| **PAM/PFS** | 3′, G-rich | 5′, T-rich | 3′, non-G-PFS | **PAMless** |
| **Substrate** | dsDNA | ssDNA, dsDNA | ssRNA only | **dsDNA, ssDNA, ssRNA** |
| **Cleavage Pattern** | Blunt | Staggered | Near U or A | Staggered |
| **Cleavage (*cis*/*trans*)** | cis | cis, trans (ssDNA) | cis, trans (RNA) | **cis/trans (dsDNA, ssDNA, ssRNA)** |
| **Guide-target duplex length** | 20 bp | 20 bp | 24 bp | 20-23 bp |

**Supplementary Table 2. crRNA sequences for *in vitro* experiments.**

| Name | Sequence |
|---|---|
| **crRNA Zero-Blunt TOPO plasmid** | AATTTCTACTGTTGTAGATTCCTAAGGCGTTACCCCAAT |
| **crRNA M13 phage** | AATTTCTACTGTTGTAGATAACGAACCACCAGCAGAAGA |
| **crRNA ssRNA** | AATTTCTACTGTTGTAGATTCCTAAGGCGTTACCCCAAT |
| **crRNA FnCas12a (Blunt TOPO plasmid)** | AATTTCTACTGTTGTAGATTCCTAAGGCGTTACCCCAAT |
| **crRNA FnCas12a (pUC18)** | AATTTCTACTGTTGTAGATCCAGTCACGACGTTGTAAAACGA |
| **crRNA SpCas9** | GTCCTAAGGCGTTACCCCAAGTTTTAGAGCTA |
| **crRNA CasΦ** | CAACGATTGCCCCTCACGAGGGGACTCCTAAGGCGTTACCCCAAT |
| **crRNA LbCas12a** | AATTTCTACTAAGTGTAGATCCAGTCACGACGTTGTAAAACGA |
| **crRNA AsCas12a** | TAATTTCTACTCTTGTAGATCCAGTCACGACGTTGTAAAACGA |
| **crRNA PAM** | AATTTCTACTGTTGTAGATGAGGGGTACCGAGCTCGAATTCG |

**Supplementary Table 3. Target sequences for targeted cleavage experiments.**

| Substrate | Sequence |
|---|---|
| **Zero-Blunt TOPO plasmid** | Commercial |
| **pUC18** | Commercial |
| **ssDNA M13 phage** | Commercial |
| **ssRNA** | GGGCCAGTGAATTCGAGCTCGGTACCCGGGGATCCT<br>CTAGAAATATGGGCTGTCCGATCGTATAACAGGATT<br>CCGCAATGGGGTTACCGCTTAAGCATTAGGGGAGCT<br>CGATTACTTGGTAGAACAGCAATCTACTCGACCTGC<br>AGGCATGCAAGCTTGGCGTAATCATGTGTTTATCCGC<br>TCACAATTCCACACAACATACGAGCCGGAAGCATAA<br>AGCGTAATCATGGTCATAGCTGTTTCCTG |

**Supplementary Table 4. Activator sequences (CIS-substrates) for non-targeted cleavage experiments.**

| Substrate | Sequence |
|---|---|
| dsDNA | GGATCCTAATACGACTCACTATAGGGACAGGCTAGCATATTGTCCTA<br>AGGCGTTACCCCAATGGCGAATTCGTAATCCCCTCGAG |
| ssDNA | CTCGAGGGGATTACGAATTCGCCATTGGGGTAACGCCTTAGGACAAT<br>ATGCTAGCCTGTCCCTATAGTGAGTCGTATTAGGATCC |
| ssRNA | CTCGAGGGGATTACGAATTCGCCATTGGGGTAACGCCTTAGGACA<br>ATATGCTAGCCTGTCGG |

**Supplementary Table 5. *Trans* substrates for non-targeted cleavage experiments.**

| Substrate | Sequence |
|---|---|
| dsDNA | pUC18 plasmid (Commercial) |
| ssDNA | M13 phage (Commercial) |
| ssRNA | GGCCAGUGAAUUCGAGCUCGGUACCCGGGGAUCCUCUAGAAAUAU<br>GGAUUACUUGGUAGAACAGCAAUCUACUCGACCUGCAGGCAUGCA<br>AGCUUGGCGUAAUCAUGGUCAUAGCUGUUUCCUGUGUUUAUCCGC<br>UCACAAUUCCACACAACAUACGAGCCGGAAGCAUAAAG |

**Supplementary Table 6. PAM library cloned in pUC18.** Target sequence is indicated in blue and random PAM in green.

| Sequence |
|---|
| tcgcgcgtttcggtgatgacggtgaaaacctctgacacatgcagctcccggagacggtcacagcttgtctgtaagcggatgccgg<br>GAGCAGACAAGCCCGTCAGGGCGCGTCAGCGGGTGTTGGCGGGTGTCGGGGC<br>TGGCTTAACTATGCGGCATCAGAGCAGATTGTACTGAGAGTGCACCATATGCG<br>GTGTGAAATACCGCACAGATGCGTAAGGAGAAAATACCGCATCAGGCGCCATT<br>CGCCATTCAGGCTGCGCAACTGTTGGGAAGGGCGATCGGTGCGGGCCTCTTCG<br>CTATTACGCCAGCTGGCGAAAGGGGGATGTGCTGCAAGGCGATTAAGTTGGGT<br>AACGCCAGGGTTTTCCCAGTCACGACGTTGTAAAACGACGGCCAGTGCCAAGC<br>TTGCATGCCTGCAGGTCGACTCTAGAGGATCCAGCAACAACGGTCGGCCACA<br>CCTTCCATTGTCGTGGCCACGCTCGGATTACACGGCAGAGGTGCTTGTGTTCCG<br>ACAGGCTAGCATATTGTCCTAAGGCGTTACCCCAANNNNNNNGGTACCGAGCT<br>CGAATTCGTAATCATGGTCATAGCTGTTTCCTGTGTGAAATTGTTATCCGCTCA<br>CAATTCCACACAACATACGAGCCGGAAGCATAAAGTGTAAAGCCTGGGGTGCC<br>TAATGAGTGAGCTAACTCACATTAATTGCGTTGCGCTCACTGCCCGCTTTCCAG<br>TCGGGAAACCTGTCGTGCCAGCTGCATTAATGAATCGGCCAACGCGCGGGGAG<br>AGGCGGTTTGCGTATTGGGCGCTCTTCCGCTTCCTCGCTCACTGACTCGCTGCG<br>CTCGGTCGTTCGGCTGCGGCGAGCGGTATCAGCTCACTCAAAGGCGGTAATAC<br>GGTTATCCACAGAATCAGGGGATAACGCAGGAAAGAACATGTGAGCAAAAGG<br>CCAGCAAAAGGCCAGGAACCGTAAAAAGGCCGCGTTGCTGGCGTTTTTCCATA<br>GGCTCCGCCCCCCTGACGAGCATCACAAAAATCGACGCTCAAGTCAGAGGTGG<br>CGAAACCCGACAGGACTATAAAGATACCAGGCGTTTCCCCCTGGAAGCTCCCT<br>CGTGCGCTCTCCTGTTCCGACCCTGCCGCTTACCGGATACCTGTCCGCCTTTCTC<br>CCTTCGGGAAGCGTGGCGCTTTCTCAAAGCTCACGCTGTAGGTATCTCAGTTCG<br>GTGTAGGTCGTTCGCTCCAAGCTGGGCTGTGTGCACGAACCCCCCGTTCAGCCC<br>GACCGCTGCGCCTTATCCGGTAACTATCGTCTTGAGTCCAACCCGGTAAGACA<br>CGACTTATCGCCACTGGCAGCAGCCACTGGTAACAGGATTAGCAGAGCGAGGT |

| |
|---|
| ATGTAGGCGGTGCTACAGAGTTCTTGAAGTGGTGGCCTAACTACGGCTACACT<br>AGAAGAACAGTATTTGGTATCTGCGCTCTGCTGAAGCCAGTTACCTTCGGAAA<br>AAGAGTTGGTAGCTCTTGATCCGGCAAACAAACCACCGCTGGTAGCGGTGGTT<br>TTTTTGTTTGCAAGCAGCAGATTACGCGCAGAAAAAAAGGATCTCAAGAAGAT<br>CCTTTGATCTTTTCTACGGGGTCTGACGCTCAGTGGAACGAAAACTCACGTTAA<br>GGGATTTTGGTCATGAGATTATCAAAAAGGATCTTCACCTAGATCCTTTTAAAT<br>TAAAAATGAAGTTTTAAATCAATCTAAAGTATATATGAGTAAACTTGGTCTGA<br>CAGTTACCAATGCTTAATCAGTGAGGCACCTATCTCAGCGATCTGTCTATTTCG<br>TTCATCCATAGTTGCCTGACTCCCCGTCGTGTAGATAACTACGATACGGGAGGG<br>CTTACCATCTGGCCCCAGTGCTGCAATGATACCGCGAGACCCACGCTCACCGG<br>CTCCAGATTTATCAGCAATAAACCAGCCAGCCGGAAGGGCCGAGCGCAGAAG<br>TGGTCCTGCAACTTTATCCGCCTCCATCCAGTCTATTAATTGTTGCCGGGAAGC<br>TAGAGTAAGTAGTTCGCCAGTTAATAGTTTGCGCAACGTTGTTGCCATTGCTAC<br>AGGCATCGTGGTGTCACGCTCGTCGTTTGGTATGGCTTCATTCAGCTCCGGTTC<br>CCAACGATCAAGGCGAGTTACATGATCCCCCATGTTGTGCAAAAAAGCGGTTA<br>GCTCCTTCGGTCCTCCGATCGTTGTCAGAAGTAAGTTGGCCGCAGTGTTATCAC<br>TCATGGTTATGGCAGCACTGCATAATTCTCTTACTGTCATGCCATCCGTAAGAT<br>GCTTTTCTGTGACTGGTGAGTACTCAACCAAGTCATTCTGAGAATAGTGTATGC<br>GGCGACCGAGTTGCTCTTGCCCGGCGTCAATACGGGATAATACCGCGCCACAT<br>AGCAGAACTTTAAAAGTGCTCATCATTGGAAAACGTTCTTCGGGGCGAAAACT<br>CTCAAGGATCTTACCGCTGTTGAGATCCAGTTCGATGTAACCCACTCGTGCACC<br>CAACTGATCTTCAGCATCTTTTACTTTCACCAGCGTTTCTGGGTGAGCAAAAAC<br>AGGAAGGCAAAATGCCGCAAAAAAGGGAATAAGGGCGACACGGAAATGTTGA<br>ATACTCATACTCTTCCTTTTTCAATATTATTGAAGCATTTATCAGGGTTATTGTC<br>TCATGAGCGGATACATATTTGAATGTATTTAGAAAAATAAACAAATAGGGGTT<br>CCGCGCACATTTCCCCGAAAAGTGCCACCTGACGTCTAAGAAACCATTATTAT<br>CATGACATTAACCTATAAAAATAGGCGTATCACGAGGCCCTTTCGTC |

**Supplementary Table 7. Primers for PAM determination assay.**

| Name | Sequence |
|---|---|
| FW library | AATAGGCGTATCACGAGGC |
| RV library | AGCGAGTCAGTGAGCGAG |

**Supplementary Table 8. *In vitro* cleavage PAM sequences.** Target sequence indicated in blue and PAM sequence in green.

| PAM | Sequence |
|---|---|
| **TTG** | TGTGAAATACCGCACAGATGCGTAAGGAGAAAATACCGCATC<br>AGGCGCCATTCGCCATTCAGGCTGCGCAACTGTTGGGAAGGG<br>CGATCGGTGCGGGCCTCTTCGCTATTACGCCAGCTGGCGAAA<br>GGGGGATGTGCTGCAAGGCGATTAAGTTGGGTAACGCCAGGG<br>TTTTCCCAGTCACGACGTTGTAAAACGACGGCCAGTGCCAAG<br>CTTGCATGCCTGCAGGTCGACTCTAGAGGATCCAGCAACAA<br>CGGTCGGCCACACCTTCCATTGTCGTGGCCACGCTCGGATTAC<br>ACGGCAGAGGTGCTTGTGTTCCGACAGGCTAGCATATTGTCCT<br>AAGGCGTTACCCCAATTGGAGGGGTACCGAGCTCGAATTCGT<br>AATCATGGTCATAGCTGTTTCCTGTGTGAAATTGTTATCCGCT |

| | |
|---|---|
| | CACAATTCCACACAACATACGAGCCGGAAGCATAAAGTGTAA<br>AGCCTGGGGTGCCTAATGAGTGA |
| **TTT** | TGTGAAATACCGCACAGATGCGTAAGGAGAAAATACCGCATC<br>AGGCGCCATTCGCCATTCAGGCTGCGCAACTGTTGGGAAGGG<br>CGATCGGTGCGGGCCTCTTCGCTATTACGCCAGCTGGCGAAA<br>GGGGGATGTGCTGCAAGGCGATTAAGTTGGGTAACGCCAGGG<br>TTTTCCCAGTCACGACGTTGTAAAACGACGGCCAGTGCCAAG<br>CTTGCATGCCTGCAGGTCGACTCTAGAGGGATCCAGCAACAA<br>CGGTCGGCCACACCTTCCATTGTCGTGGCCACGCTCGGATTAC<br>ACGGCAGAGGTGCTTGTGTTCCGACAGGCTAGCATATTGTCCT<br>AAGGCGTTACCCCAATTTGAGGGGTACCGAGCTCGAATTCGT<br>AATCATGGTCATAGCTGTTTCCTGTGTGAAATTGTTATCCGCT<br>CACAATTCCACACAACATACGAGCCGGAAGCATAAAGTGTAA<br>AGCCTGGGGTGCCTAATGAGTGA |
| **CCC** | TGTGAAATACCGCACAGATGCGTAAGGAGAAAATACCGCATC<br>AGGCGCCATTCGCCATTCAGGCTGCGCAACTGTTGGGAAGGG<br>CGATCGGTGCGGGCCTCTTCGCTATTACGCCAGCTGGCGAAA<br>GGGGGATGTGCTGCAAGGCGATTAAGTTGGGTAACGCCAGGG<br>TTTTCCCAGTCACGACGTTGTAAAACGACGGCCAGTGCCAAG<br>CTTGCATGCCTGCAGGTCGACTCTAGAGGGATCCAGCAACAA<br>CGGTCGGCCACACCTTCCATTGTCGTGGCCACGCTCGGATTAC<br>ACGGCAGAGGTGCTTGTGTTCCGACAGGCTAGCATATTGTCCT<br>AAGGCGTTACCCCAACCCGAGGGGTACCGAGCTCGAATTCGT<br>AATCATGGTCATAGCTGTTTCCTGTGTGAAATTGTTATCCGCT<br>CACAATTCCACACAACATACGAGCCGGAAGCATAAAGTGTAA<br>AGCCTGGGGTGCCTAATGAGTGA |
| **TCA** | TGTGAAATACCGCACAGATGCGTAAGGAGAAAATACCGCATC<br>AGGCGCCATTCGCCATTCAGGCTGCGCAACTGTTGGGAAGGG<br>CGATCGGTGCGGGCCTCTTCGCTATTACGCCAGCTGGCGAAA<br>GGGGGATGTGCTGCAAGGCGATTAAGTTGGGTAACGCCAGGG<br>TTTTCCCAGTCACGACGTTGTAAAACGACGGCCAGTGCCAAG<br>CTTGCATGCCTGCAGGTCGACTCTAGAGGGATCCAGCAACAA<br>CGGTCGGCCACACCTTCCATTGTCGTGGCCACGCTCGGATTAC<br>ACGGCAGAGGTGCTTGTGTTCCGACAGGCTAGCATATTGTCCT<br>AAGGCGTTACCCCAATCAGAGGGGTACCGAGCTCGAATTCGT<br>AATCATGGTCATAGCTGTTTCCTGTGTGAAATTGTTATCCGCT<br>CACAATTCCACACAACATACGAGCCGGAAGCATAAAGTGTAA<br>AGCCTGGGGTGCCTAATGAGTGA |
| **TGG** | TGTGAAATACCGCACAGATGCGTAAGGAGAAAATACCGCATC<br>AGGCGCCATTCGCCATTCAGGCTGCGCAACTGTTGGGAAGGG<br>CGATCGGTGCGGGCCTCTTCGCTATTACGCCAGCTGGCGAAA<br>GGGGGATGTGCTGCAAGGCGATTAAGTTGGGTAACGCCAGGG<br>TTTTCCCAGTCACGACGTTGTAAAACGACGGCCAGTGCCAAG<br>CTTGCATGCCTGCAGGTCGACTCTAGAGGGATCCAGCAACAA<br>CGGTCGGCCACACCTTCCATTGTCGTGGCCACGCTCGGATTAC<br>ACGGCAGAGGTGCTTGTGTTCCGACAGGCTAGCATATTGTCCT<br>AAGGCGTTACCCCAATGGGAGGGGTACCGAGCTCGAATTCGT<br>AATCATGGTCATAGCTGTTTCCTGTGTGAAATTGTTATCCGCT<br>CACAATTCCACACAACATACGAGCCGGAAGCATAAAGTGTAA<br>AGCCTGGGGTGCCTAATGAGTGA |

#### Supplementary Table 9. Sequences of pre-crRNA transcript.

| Substrate | Sequence |
|---|---|
| **pre-crRNA Control** | TACGCCAGCTGGCGAAAGGGGGATGTGCTGCAAGGCGATTAAGTTGGGTAACGCCAGGGTTTTCCCAGTCACGACGTTGTAAAACGACGGCCAGTGAATTCGAGCTCGGTACCCGGGGAGAAGTCATTTAATAAGGCCACTGTTAAAAAGCTTGGCGTAATCATGGTCATAGCTGTTTCCTGTGTGAAATTGTTATCCGCTCACAATTCCACACAACATACGAGCCGGAAGCATAAAGTGTAAAGCCTGGGGTGCCTA ATGAGTGAGCTAACTCACATTAATTGCGTT |
| **pre-crRNA** | GGGGGTCTTTTTTTGCTGATTTAGGCAAAAACGGGTCTAAGAACTTTAAATAATTTCTACTGTTGTAGATGAGAAGTCATTTAATAAGGCCACTGTTAAAAGTCTAAGAACTTTAAATAATTTCTACTGTTGTAGATGCTACTATTCCTGTGCCTTCAGATAATTCAGTCTAAGAACTTTAAATAATTTCTACTGTTGTAGATGTCTAGAGCCTTTTGTATTAGTAGCCGGTCTAAGAACTTTAAATAATTTCTACTGTTGTAGATTAGCGATTTATGAAGGTCATTTTTTTGTCTAGCTTTAATGCGGTAGTTTATCACAGTTAAATTGCTAACG |

#### Supplementary Table 10. Sequences for plasmid clearance assay in *E. coli*.

| Name | Sequence |
|---|---|
| Target site | CCAGTCACGACGTTGTAAAACGA |
| Plasmid | pAF (A modified version of pBAD33 plasmid containing a subcloned LacZ gene) |

#### Supplementary Table 11. Loci sequences and primers used for endogenous gene editing in human HEK293T cells.

| Locus | PAM | crRNA |
|---|---|---|
| DNMT1 | TTTC | AATTTCTACTGTTGTAGATCTGATGGTCCATGTCTGTTACTC |
| EMX1 | TTTG | AATTTCTACTGTTGTAGATTCCTCCGGTTCTGGAACCACACC |
| AAVS1 | TTTG | AATTTCTACTGTTGTAGATCTTACGATGGAGCCAGAGAGGAT |

| Name | Primer sequence |
|---|---|
| DNMT1 FW | CTGGGACTCAGGCGGGTCAC |
| DNMT1 RV | CCTCACACAACAGCTTCATGTCAGC |
| EMX1 FW | CCATCCCCTTCTGTGAATGT |
| EMX1 RV | GGAGATTGGAGACACGGAGA |
| AAVS1 FW | GGGCTGGCTACTGGCCTTAT |
| AAVS1 RV | TATCTGTCCCCTCCACCCCA |

### Supplementary Table 12. crRNAs and oligos used for cryo-EM.

| | **Sequence** |
|---|---|
| crRNA | AATTTCTACTGTTGTAGATTCCTAAGGCGTTACCCCAAT |
| Cis_DNA FW | CTAGCATATTTATCCTAAGGCGTTACCCCAATGGGAGGG |
| Cis_DNA RV | CCCTCCCATTGGGGTAACGCCTTAGGATAAATATGCTAG |
| Trans_DNA FW | AACTEFZFZEFOFFZZEOEOEZAGCA |
| Trans_DNA RV | TGCTFOEOEOFFZZEZOFZFZOAGTT |

**Being: F=A, O=C, E=G and Z=T.*

### Supplementary Note 1. Structure of apo α-synCas

#### α-synCas apo state structure determination

The cryo-EM structure of apo α-synCas was obtained fortuitously. We initially aimed to determine the cryo-EM structure of the binary complex of α-synCas bound to crRNA. However, despite several attempts we failed at producing a stable binary complex that survived vitrification and ended up with the apo form of α-synCas, which upon data processing (Supplementary Fig. 3) yielded a 3.3Å resolution 3D reconstruction (Supplementary Fig. 2; Extended Table 1). The structure of apo α-synCas without crRNA displays a ring-like architecture distinct from the oval "sea conch" shape of the ternary complex (Supplementary Fig. 4). The REC1 and REC2 domains in the apo state occupy different positions relative to the ternary complex, and the charged central channel between the recognition and nuclease lobes, through which the crRNA-target DNA heteroduplex threads, is not formed. The REC2 domain closes the ring by connecting with the RuvC domain, in a way that hinders access to the nuclease active site (Supplementary Fig. 4). More specifically, helical residues 378-390 of REC2 contact the lid loop residues 970-980 of the RuvC domain, through electrostatic and π-stacking interactions (D379-R977, D384-R975, W379- F978, K383-F973).

### Supplementary Note 2. Detailed Cryo-EM materials and methods

***Specimen preparation and cryo-EM data collection***. Purified α-synCas following 3C elution from affinity chromatography was mixed with crRNA in a 1:1.2 α-synCas:crRNA molar ratio (Supplementary Table 12), and after 15 minutes of incubation the mixture was run over a Superose 6 GL 10/300 column to remove excess crRNA by size exclusion chromatography (SEC). The SEC fractions of the binary complex were pooled down and concentrated to 2.5 mg/ml. One aliquot of this preparation was used for cryo-EM, however, the binary complex did not survive vitrification, as the enzyme unbound the crRNA, and we could only determine the structure of apo α-synCas. Another aliquot of the SEC purified binary complex was mixed with target dsDNA in a 1:1.2:3 molar ratio of α-synCas:crRNA:dsDNA (Supplementary Table 12). This sample yielded a stable ternary complex that withstood vitrification. To the ternary complex mixture we added collateral dsDNA in a 1:1.2:3:3 molar ratio of α-synCas:crRNA:target-dsDNA:collateral-dsDNA (Supplementary Table 12). We added up to 0.05% CHAPS to all specimens before vitrification to avoid preferred orientations. For cryo-EM, 4 ul aliquots of the different α-synCas complexes were adsorbed onto glow-discharged Quantifoil R 1.2/1.3 300 mesh grids (Quantifoil), and vitrified in liquid ethane with a Leica EM GP2 cryoplunger (Leica) using front-side blotting for 2 s at 95% humidity. The specimens were imaged in house using a 300 kV Krios G4 (ThermoScientific) equipped with a BioContinuum/K3 camera (Gatan) operating in counting mode at a calibrated 0.8238 Å/pix. Employing a 1-1.6 μm underfocus range we recorded three movies per hole with a total accumulated dose of 50 e−/Å2 over 50 frames. Movies were recorded automatically using EPU 2 (ThermoScientific) with Aberration-free image shift (AFIS) and Fringe-free imaging (FFI).

***Cryo-EM data processing and 3D reconstruction***. Processing of image data followed a processing strategy based on methodologies implemented in cryoSPARC[1] . Initial frame alignment and contrast transfer function (CTF) estimation was performed using cryoSPARC live[1]. Movies selected according to CTF and total motion were chosen for particle picking. Blob picker was used for picking particles that were cleaned in 2D-classifications to produce templates for template particle picking and to train the neural-network picker Topaz[2]. For the ternary and quaternary complexes, Topaz picked particles were used to produce initial models for heterogeneous refinement using Ab-initio reconstruction (Supplementary Fig. 1). Further 3D classifications of the best class particles followed by non-uniform refinements yielded 3D reconstructions of the ternary and quaternary complexes at 3.14 Å and 3.08 Å resolution respectively (Supplementary Fig. 3).

For apo α-synCas (Supplementary Fig. 2), a subset of micrographs was used to generate initial models by Ab-initio reconstruction. Particles picked using template and Topaz were cleaned using 2D-classifications and merged, and duplicates were removed. Heterogeneous refinement was performed using the generated initial models, and the best class was further classified in heterogeneous refinements followed by non-uniform refinement, yielding a final 3D reconstruction of the apo state at 3.3 Å resolution (Supplementary Fig. 3). To improve the cryo-EM density of the flexible REC1 domain a 3D variability analysis (3DVA)[3] was performed followed by a heterogeneous refinement. The particles of the best class were subjected to non-uniform refinement yielding a 3D reconstruction at 3.75 Å resolution showing a more complete density for REC1 domain. In addition, the second class of the first heterogenous refinement was further classified and refined obtaining a noisy 3D reconstruction that lacks density for the REC domains.

*Model building*. For initial atomic model building from the cryo-EM density maps we employed automated model building with ModelAngelo[4]. In the case of apo state, the model was of high quality for most of the structure, however the REC1 domain was further completed by manual intervention in Coot[5] using the REC1-improved cryo-EM map. The final model lacks residues 128-148 (REC1), 392-412 and 464-490 (REC2), 661-676 (PI), 796-834 and 754-761 (Wed.), and 902-939 (RuvC and BH), due to their respective absence of density in the cryo-EM map. For the ternary complex the initial model was also completed by manual intervention in Coot[5] , except for residues 215-234 and 258-268 (REC1), and 302-313 (REC1-REC2 linker), and nucleotides NTS 14-27, TS 18-27 and crRNA 14-20. The REC2 domain was fitted as a rigid body. In the case of the quaternary complex, the atomic model of the ternary complex was used as an initial model. The model was improved in Coot, lacking due to poor density the residues 216-230 and 258-268 (REC1) and 302-313 (REC1-REC2 linker), and nucleotides NTS 5-27, TS 18-27 and crRNA 14-20, that do not show density in the cryo-EM map. To improve the fit and to optimize stereochemistry real space refinements with secondary structure and geometry restraints were performed using Phenix[6]. Finally, an ideal four-nucleotide dsDNA was generated in Coot and rigid body fitted into the unassigned cryo-EM density of the quaternary complex cryo-EM map for visualization purposes.

*Figure preparation*. All the figures were prepared using CorelDRAW and Adobe Illustrator. Cryo-EM maps and structures were visualized, and their images generated using ChimeraX[7].
